# Rational design of a SOCS1-edited tumor infiltrating lymphocyte therapy for solid tumors using CRISPR/Cas9 screens

**DOI:** 10.1101/2023.09.05.555798

**Authors:** Michael R. Schlabach, Sharon Lin, Zachary Collester, Christopher Wrocklage, Sol Shenker, Conor Calnan, Tianlei Xu, Hugh Gannon, Leila Williams, Frank Thompson, Paul Dunbar, Robert A. LaMothe, Tracy E. Garrett, Nick Colletti, Anja F. Hohmann, Noah Tubo, Caroline Bullock, Isabelle Le Mercier, Katri Sofjan, Jason J. Merkin, Sean Keegan, Gregory V. Kryukov, Caroline Dugopolski, Frank Stegmeier, Karrie Wong, Fiona A. Sharp, Louise Cadzow, Micah J. Benson

**Affiliations:** KSQ Therapeutics, 4 Maguire Road, Lexington, MA 02421

## Abstract

Cell therapies such as Tumor Infiltrating Lymphocyte (TIL) therapy have shown promise in the treatment of patients with refractory solid tumors, with improvement in response rates and durability of responses nevertheless sought. To identify targets capable of enhancing the anti-tumor activity of T cell therapies, large-scale in vitro and in vivo CRISPR/Cas9 screens were performed, with the suppressor of cytokine signaling 1 (SOCS1) gene identified as a top T cell-enhancing target. In murine CD8 T cell therapy models, SOCS1 served as a critical checkpoint in restraining the accumulation of T central memory cells in lymphoid organs as well as intermediate (Tex^int^) and effector (Tex^eff^) exhausted T cell subsets derived from progenitor exhausted T cell (Tex^prog^) cells in tumors. A comprehensive CRISPR tiling screen of the *SOCS1* coding region identified sgRNAs targeting the SH2 domain of SOCS1 as the most potent, with a sgRNA with minimal off-target cut sites used to manufacture KSQ-001, an engineered TIL therapy with SOCS1 inactivated by CRISPR/Cas9. KSQ-001 possessed increased responsiveness to cytokine signals and enhanced in vivo anti-tumor function in mouse models. These data demonstrate the use of CRISPR/Cas9 screens in the rational design of T cell therapies.

## Introduction

Cell therapies using T lymphocytes possessing anti-tumor specificity have demonstrated robust clinical activity in hematological malignancies(1–4), with more limited success observed in solid tumor indications(5–7). The reduced effectiveness of T cell therapies in solid tumors is thought to be due, in part, to the immunosuppressive nature of solid tumor and its blunting impact on the function of transferred T cells. Despite this challenge of tumor-driven immunosuppression, T cell therapies using Tumor Infiltrating Lymphocytes (TIL) derived from a patient’s tumor have demonstrated promise in certain clinical settings, such as in refractory metastatic melanoma, with objective responses observed in patients who have failed multiple rounds of therapy, including with immune-checkpoint inhibitors(8–11). However, the majority of melanoma patients receiving TIL ultimately progress, and fewer objective responses have been observed in other solid tumor indications, such as in cervical(12), ovarian(13) and NSCLC(14), where the immunosuppressive TME barrier may be more stringent then in melanoma. Strategies are thus sought to enhance the anti-tumor potency, accumulation, persistence, and memory formation of TIL and other T cell therapies as enhancement in these functional attributes are predicted to directly translate to increased durable objective responses. It is currently unclear which T cell pathways are most impactful to target in enhancing the anti-tumor function of T cell therapies against solid tumors.

Functional screens constitute a powerful approach to discover novel biology in disease-relevant mammalian models. The ability of the Clustered Regularly Interspaced Short Palindromic Repeats (CRISPR) / Cas9 gene editing system to precisely inactivate a target gene at the level of genomic DNA has enabled the large-scale interrogation of gene function by use of pooled loss-of-function screens(15–17). This approach has been notably used for the discovery of genetic vulnerabilities and therapeutic targets in cancer types(18) and to immune cell types in various settings, including the mapping of innate cells in response to pro-inflammatory stimuli(19, 20), pathways involved in regulating T cell function (21–24) and regulatory T cell function(25, 26). CRISPR/Cas9 functional genomic screens thus afford the opportunity to comprehensively map the biology of primary T cells by querying gene targets driving pre-defined functions.

In this study, we used large-scale CRISPR/Cas9 screens to identify top T cell therapy enhancing targets, with SOCS1 identified as a top hit. SOCS1 serves as a negative regulator of cytokine signaling in T cells, including IL-2, IL-12 and IL-15, with these cytokines known to influence the survival, differentiation and function of T cells (27, 28). SOCS1 was found to be a checkpoint in transferred CD8 T cells restraining the accumulation of CD44^+^CD62L^+^ T central memory cells within peripheral lymphoid organs as well as in the differentiation of Slamf6^+^CD39^-^PD-1^med^ progenitor T exhausted (Tex^prog^) cells into Slamf6^-^CD39^+^PD-1^hi^ intermediate (Tex^int^) and effector (Tex^eff^) subsets within tumors. Based on these insights, we used a SOCS1 protein domain CRISPR/Cas9 tiling screen to develop KSQ-001, a engineered TIL (eTIL®) therapy with inactivation of the *SOCS1* gene via CRISPR/Cas9 editing that demonstrated enhanced production of IFNγ, increased responsiveness to cytokine signals and increased anti-tumor activity in a solid tumor TIL model. The use of CRISPR/Cas9 functional genomic screens in the rational design of engineered TIL is hereby demonstrated.

## Results

### Functional CRISPR/Cas9 screens identify SOCS1 as a top target constraining in vitro expansion of TIL and in vivo infiltration of transferred CD8 T cells in syngeneic solid tumor models

To identify targets that can enhance TIL function, we performed large-scale *in vitro* and *in vivo* CRISPR/Cas9 screens. The manufacture of TIL for therapeutic use occurs in a multi-step process involving surgical resection of tumor, extraction of TIL from tumor fragments in the presence of IL-2 in a step known as a ‘pre-REP,’ followed by expansion of TIL in the presence of IL-2, irradiated PBMC feeders and agonistic anti-CD3 OKT3 antibody in the Rapid Expansion Phase, or REP (29–32). As attaining the appropriate dose level of TIL can be a challenge, especially with heavily pre-treated patients, a CRISPR was performed on TIL derived from single a metastatic melanoma donor to identify gene targets enhancing expansion under REP conditions. A barcoded sgRNA library encompassing a curated list of 5137 genes involved in T cell function, all predicted cell surface receptors (CSRs), and genes found to be expressed within immune cells and peripheral blood was introduced by lentiviral transduction into pre-REP TIL together with Cas9. Engineered sgRNA Lib^+^ TIL were then expanded under REP conditions (Figure 1A). In this screen, SOCS1 was identified as the top hit driving the expansion of TIL in REP conditions (FDR adjusted p-value: 4.03x10^-4^) (Figure 1B, Table S1). Confirming screen robustness, we observed recovery of 85% library sgRNAs as well as depletion of sgRNAs targeting essential genes, including strong depletion of RPL10A, as well as multiple members of the IL-2 receptor signaling complex, including IL-2RG, STAT5A, JAK3 and IL-2RB (Supplemental Data 1, Figure 1b), consistent with the dependency of TIL on IL-2 survival signals during a REP.

**Figure 1:**
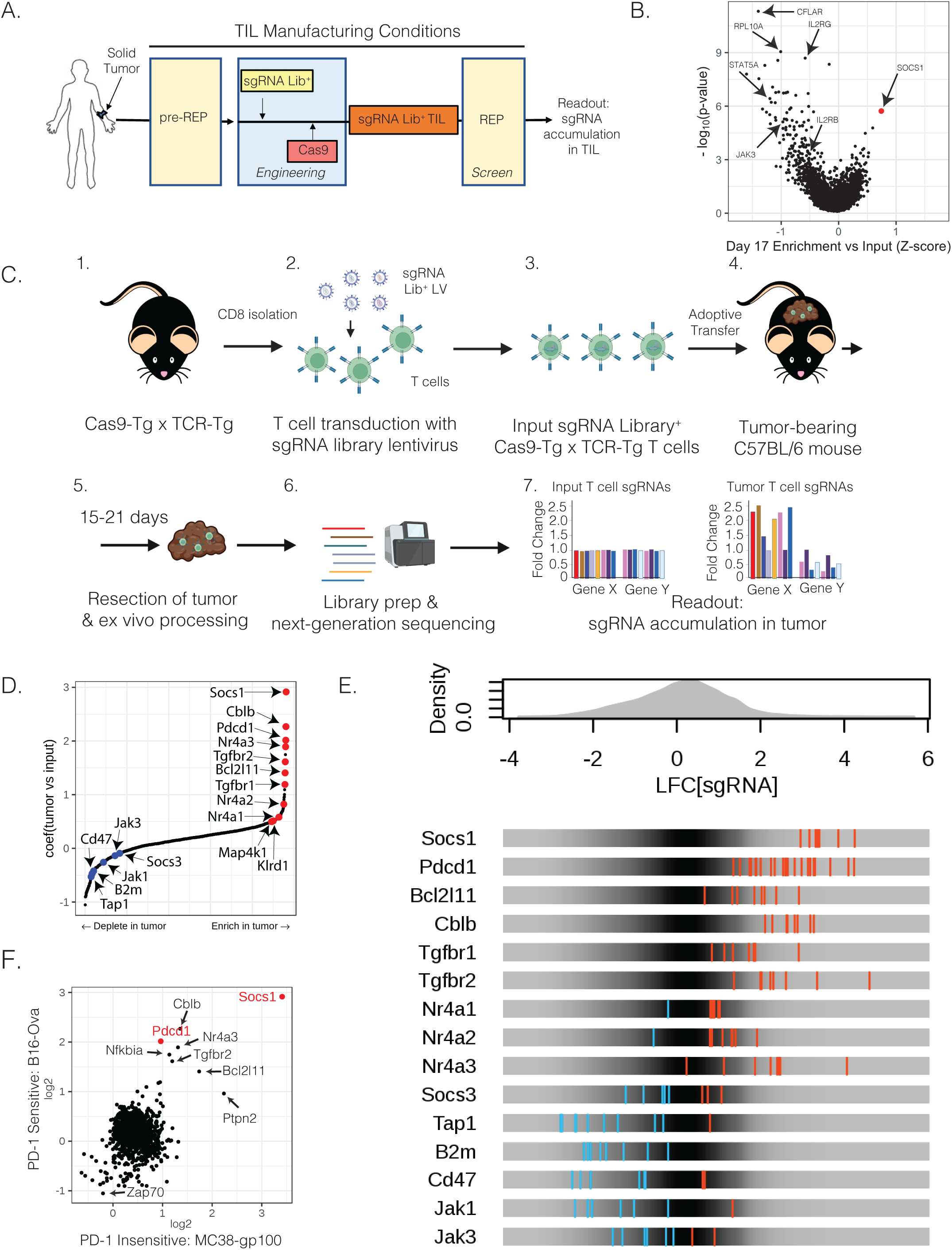
Functional CRISPR/Cas9 screens identify SOCS1 as a top target constraining in vitro expansion of TIL and in vivo infiltration of transferred CD8 T cells in syngeneic solid tumor models. **(A)** Experimental schematic depicting a CRISPR screen performed on the in vitro expansion of human melanoma TIL under conditions used to manufacture for therapeutic use. Following pre-REP, a sgRNA Library was introduced by lentiviral transduction with TIL then engineered by introduction of Cas9 by electroporation. A REP was then initiated in the presence of allogeneic iPBMC feeders, OKT3 and IL-2. The distribution of sgRNAs was compared prior to and following the REP to identify targets enhancing TIL accumulation. **(B)** SOCS1 is the top target enhancing the accumulation of TIL under REP conditions. **(C)** Experimental schematic depicting the workflow of in vivo CRISPR screens using Cas9-Tg x TCR-Tg (OT1 or PMEL) CD8 T cells. CD8 T cells were activated and transduced to express a sgRNA Library, with Cas9-Tg x TCR-Tg T cells transferred into mice bearing 100mm^3^ tumors on the flank. Either 14- or 21-days following transfer, tumors were harvested and the sgRNA distribution of T cells within tumors analyzed and compared with the input population of T cells. **(D)** MAGeCK-MLE identifies SOCS1 as a top target enhancing OT1 T cell infiltration into B16-Ova tumors 14 days following transfer. **(E)** Enrichment or depletion patterns of sgRNAs targeting known genes by tumor OT1s in comparison to input OT1s. **(F)** MAGeCK-MLE identifies SOCS1 as a top target enhancing the infiltration of PMEL CD8 T cells 14 days after transfer into the MC38-gp100 solid tumor model found to be refractory to inhibition of PD-1.

To identify targets that enhance the ability of CD8 T cells to infiltrate, accumulate and persist in solid tumors in vivo, we crossed Cas9 transgenic mice(33) with OT1 TCR-Tg mice with the Cas9-Tg x OT1 CD8 T cells recognizing and Ovalbumin (Ova) peptide (34). We conducted a CRISPR screen in the B16-Ova model, found to be sensitive to transferred OT1 T cells in which the *Pdcd1* gene encoding PD-1 has been inactivated using CRISPR/Cas9 (Figure S1A). Two barcoded sgRNA libraries were introduced into Cas9-Tg x OT1 T cells covering a curated list of 369 T cell genes as well as 1004 predicted cell surface receptors. Following transfer of sgRNA^+^ Cas9-Tg x OT1 T cells into B16-Ova tumor-bearing mice, the distribution of sgRNAs present within the transferred cells infiltrating tumor was evaluated 14 or 21 days later (Figure 1C). We observed recovery of 97% of sgRNAs from tumor samples (Supplemental Data 2) and strong enrichment of sgRNAs targeting the *Pdcd1* gene (FDR-adjusted p-value: < 1x10^-16^). Confirming screen robustness, sgRNAs targeting genes required for T cell function were depleted from the tumor, including *Tap1, B2m, CD47, Jak1* and *Jak3* (Figure 1D-E). In addition, our screen showed enrichment of sgRNAs targeting other genes encoding negative regulators of T cell function, including *Bcl2l11*, *Cblb,* both *Tfgbr1* and *Tgfbr2, Kldr1* and all three members of the *Nr4a* family (Figure 1D-E). Notably, sgRNAs targeting the *Socs1* gene encoding SOCS1 emerged as the strongest target driving enhanced CD8 T cell infiltration within tumors (FDR adjusted p-value: < 1x10^-16^) greater than sgRNAs targeting *Pdcd1* (Figures 1D-E; Table S2).

The highest unmet medical need for the treatment of solid tumors is in the PD-1 refractory setting. We therefore performed an additional in vivo CRISPR/Cas9 screen using pmel CD8 T cells in a MC38-gp100 tumor model found to be insensitive to treatment by *Pdcd1*-inactivated pmel T cells(35) The screen again performed robustly with 82% of the sgRNAs recovered (Supplemental Data 3) and with sgRNAs targeting the *Pdcd1* gene displaying no enrichment within tumors (Figure 1F). By contrast, *Socs1* again scored as the top T cell target in this screen (FDR adjusted p-value: < 1x10^-16^) (Figure 1F). In this particular screen, other T cell targets which enriched in the OT1/B16-Ova screen did not demonstrate as strong enrichment, including *Cblb, Bcl2l11, Tgfbr2, Nr4a3* and *Nfkbia* (Figure 1F; Table S2).

Collectively, these data identify SOCS1 as a target which, upon CRISPR/Cas9-mediated inactivation, enhances the ability of TIL to expand under manufacturing conditions and to drive the infiltration and accumulation of tumor-specific CD8 T cells within solid tumors in an in vivo setting.

### Inactivation of SOCS1 by CRISPR/Cas9 in transferred CD8 T cells drives enhanced efficacy in syngeneic mouse models with durable persistence as T_cm_ cells

We next evaluated the impact of inactivating SOCS1 on the anti-tumor function and memory formation of transferred CD8 T cells. OT1 T cells were edited using CRISPR/Cas9 RNPs targeting either *Socs1* (sgSocs1), *Pdcd1* (sgPD-1), or *Olf1a* (sgOlf) as a negative control and were transferred into mice bearing established 100mm^3^ B16-Ova tumors. Editing efficiencies for target genes were 82% for sgPD-1, 92% for sgSocs1, and 80% for sgOlf. Mice receiving sgSocs1-edited OT1 T cells displayed clearance of tumors, with 10/10 mice undergoing complete rejection (CR) of tumor in comparison to the inadequate tumor control demonstrated by mice receiving sgPD-1 or sgOlf OT1s (Figure 2A). sgSocs1 OT1-treated mice undergoing a CR were re-challenged with B16-Ova tumors 76 days following initial transfer and completely rejected secondary tumor challenge (Figure 2B). The persistence and phenotype of sgSocs1 OT1s were tracked longitudinally in the peripheral blood of CR mice both immediately prior to and following B16-Ova re-challenge. Following initiation tumor rejection, sgSocs1 OT1s comprised 7% of blood CD8 T cells (Figure 2C) with a CD44^+^CD62L^+^ phenotype reflecting T central memory (T_cm_) cells (Figure 2D). Following re-challenge, sgSocs1 OT1s expanded to occupy 37% of totally blood CD8^+^ cells and displayed a CD44^+^CD62L^-^ T effector memory (T_em_) cell phenotype (Figure 2D), followed by contraction to a new baseline of 24% of total CD8 T cells which persisted out to 160 days until the termination of the experiment, and displaying a CD44^+^CD62L^+^ T_cm_ phenotype (Figure 2D).

**Figure 2:**
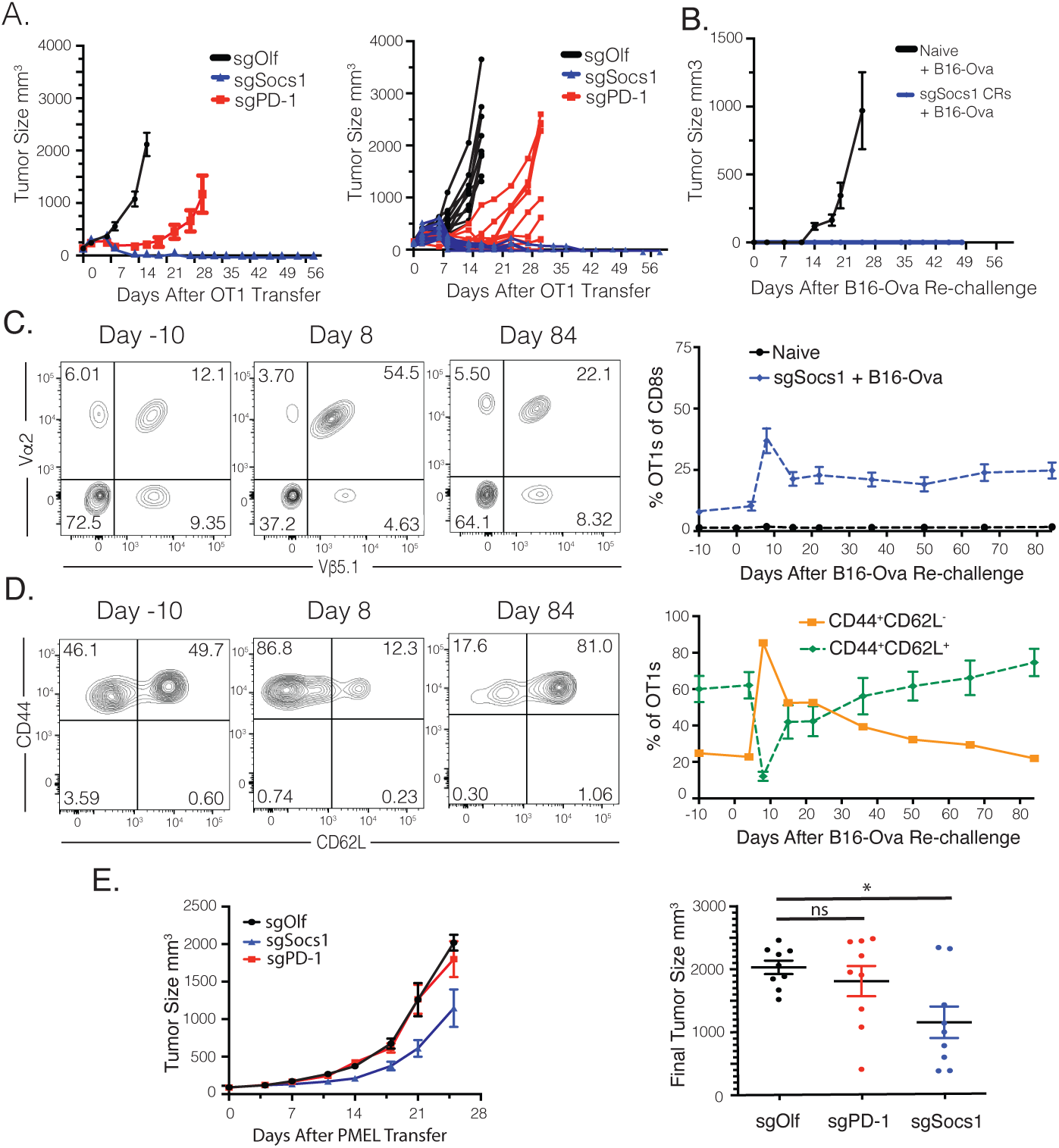
Inactivation of SOCS1 by CRISPR/Cas9 in transferred CD8 T cells drives enhanced efficacy in syngeneic mouse models with durable persistence as T_cm_ cells. C57BL/6 mice bearing 100mm^3^ B16-Ova tumors on the flank were treated with 3x10^6^ OT1 CD8 T cells engineered to inactivate either OLF1 (sgOlf), PD-1 (sgPD-1), or SOCS1 (sgSocs1). Results of statistical analysis depicted between sgOlf vs sgPD-1 and sgOlf vs sgSocs1. **(A)** Tumor growth curves of each group over time are depicted. **(B)** Mice treated with sgSocs1 OT1s undergoing complete tumor rejection were re-challenged with B16-Ova tumor cells 61 days following initial transfer, with 10 naïve mice included as controls. Tumor growth of the indicated treatment groups are depicted. **(C)** The frequency of sgSocs1 OT1s as peripheral blood CD8s prior to (Day -10) and following (Day 8 through 84) B16-Ova re-challenge. FACS plots are depicted, with OT1s defined as CD8^+^Vα2^+^Vβ5.1^+^cells. **(D)** The CD44^+^CD62L^+^ and CD44^+^CD62L^-^ phenotype of sgSocs1 OT1s in (c) prior to (Day -10) and following B16-Ova re-challenge (Day 8 and 84) was quantified by FACS and depicted. **(E)** C57BL/6 mice bearing MC38-gp100 tumors with a median size of 100mm^3^ were treated with 7x10^6^ PMEL CD8 T cells inactivated with either SOCS1 (sgSocs1), PD1 (sgPD-1) or OLF1 (sgOlf). Tumor growth curves depicted by treatment group on graph on the left, and the final tumor size in individual mice depicted on graph on the right. Results of statistical analysis depicted between sgOlf vs sgPD-1 and sgOlf vs sgSocs1. A 2-way ANOVA was used determine statistical significance between treatment groups in Figures 2a and 2e, with **** = p value < 0.0001, and ns = no significance. Figures 2A-D representative of 2 independent experiments.

To evaluate SOCS1 inactivation in transferred CD8 T cells in a syngeneic model refractory to PD-1 inactivation, sgOlf, sgSocs1 or sgPD-1 pmel cells were adoptively transferred into mice bearing established 100mm^3^ MC38-gp100 tumors. Mice receiving sgSocs1 pmels showed controlled tumor growth, with hosts receiving sgPD-1 edited or sgOlf pmels showing no impact on tumor growth (Figure 2E). To quantify how inactivation of Socs1 in T cells improves anti-tumor potency in an in vivo setting, we performed a relative potency study in which the anti-tumor activity of a dose titration of sgOlf and sgSocs1 OT1s was evaluated. A dose of 4.1x10^6^ sgSocs1 OT1s matched the anti-tumor response profile generated by 41x10^6^ sgOlf-edited OT1s, suggesting that inactivation of *Socs1* leads to an estimated ten-fold anti-tumor potency increase in OT1 T cells (Figure S2A). In the 41x10^6^ dose-level cohort where both sgOlf and sgSocs1 treatment groups showed similarly strong anti-tumor activity, sgSocs1 OT1s were present at a five-fold greater frequency within the peripheral blood in comparison to sgOlf OT1 treated mice, establishing that inactivation of SOCS1 enhances the overall accumulation of transferred T cells even in similarly responding mice (Figure S2B). sgSocs1 OT1s additionally accumulated to an enhanced degree as CD44^+^CD62L^+^ T_cm_ cells in comparison to sgOlf OT1s (Figure S2C).

To conclude, inactivation of SOCS1 in CD8 T cells leads to a robust increase in anti-tumor activity in vivo including activity in models where inactivation of PD-1 on CD8 T cells has no effect. SOCS1 inactivation led to an increase in the frequency of transferred T cells detectable within the peripheral blood, with these cells possessing a T_cm_ phenotype and remaining prominent within the host as far out as 160 days following transfer.

### Inactivation of SOCS1 in transferred CD8 T cells enhances their accumulation as CD44^+^CD62L^+^ T_cm_ cells within lymphoid organs and Slamf6^-^CD39^+^PD-1^hi^ Tex cells in tumors while depleting intratumoral Tregs

We next sought to better understand how SOCS1 inactivation impacted CD8 T cell function within tumors and lymphoid organs during the acute phase of the anti-tumor response. We evaluated the frequency and phenotype of transferred sgSocs1, sgPD-1 and sgOlf OT1 T cells 7 days following transfer, with editing efficiencies for target genes 82% for sgPD-1, 95% for sgSocs1, and 80% for sgOlf. We confirmed our initial screen observation that sgSocs1 OT1s enriched to a greater degree within tumor, with sgSocs1 OT1s comprising 53% of total CD8s versus 16% of sgOlf and 23% of sgPD-1 OT1s (Figure 3A-B). A similar enrichment of sgSocs1 OT1s was observed in the peripheral blood, spleen and tumor-draining inguinal lymph nodes (TDLN) (Figure 3B). In evaluating the phenotype of transferred OT1s, a near doubling in frequency of CD44^+^CD62L^+^ T_cm_ OT1 cells was observed in the spleen and TDLN in the sgSocs1 treatment group in comparison to sgOlf and sgPD-1 OT1s (Figure 3C). Conversely, upon evaluating OT1s infiltrating tumor, a heterogenous pattern of CD62L expression was observed with no detectable expression of CD44 (Figure 3C). Tumor-infiltrating OT1s were further assessed to discern Tex differentiation state. In the T cell exhaustion differentiation continuum, TCF-1-expressing stem-like Tex^prog^ cell subsets can be identified using Slamf6 (36). Tex^prog^ cells undergo clonal expansion and differentiation into Tex subsets specializing in effector function and terminal differentiation, Slamf6 expression is lost and CD39 expression induced, with PD-1 increasing in expression (37, 38). Tumor-infiltrating OT1s across treatment groups could be divided into Slamf6^+^CD39^-^ Tex^prog^ and Slamf6^-^CD39^+^ cells encompassing Tex^int^, Tex^eff^ and Tex^term^ subsets (Figure 3D). No impact was found on the frequencies of Slamf6^+^CD39^-^ Tex^prog^ subsets between groups (Figure 3D). However, inactivation of SOCS1 in sgSocs1 OT1s drove a marked enhancement in the accumulation of the Slamf6^-^CD39^+^ population in comparison to sgOlf and sgPD-1 treatment groups (Figure 3D). Additional phenotypic analysis confirmed expression of CD62L by the Slamf6^+^CD39^-^ population in comparison to the Slamf6^-^CD39^+^ population, and with Slamf6^-^CD39^+^ OT1s conversely expressing higher levels of PD-1 protein, apart from the sgPD-1 OT1 treatment group, which was confirmed to not express detectable levels of PD-1 protein (Figure 3E). To summarize, inactivation of SOCS1 in transferred CD8 T cells led to 1) enhanced accumulation in blood, spleen, TDLN and tumor 2) increased accumulation as CD44^+^CD62L^+^ T_cm_ cells in the spleen and TDLN 3) no impact on the frequency of Slamf^+^CD39^-^PD-1^med^ Tex^prog^ cells while 4) markedly increasing the accumulation of Slamf^-^CD39^+^PD-1^hi^ cells encompassing Tex^int^, Tex^eff^ and Tex^term^ subsets.

**Figure 3:**
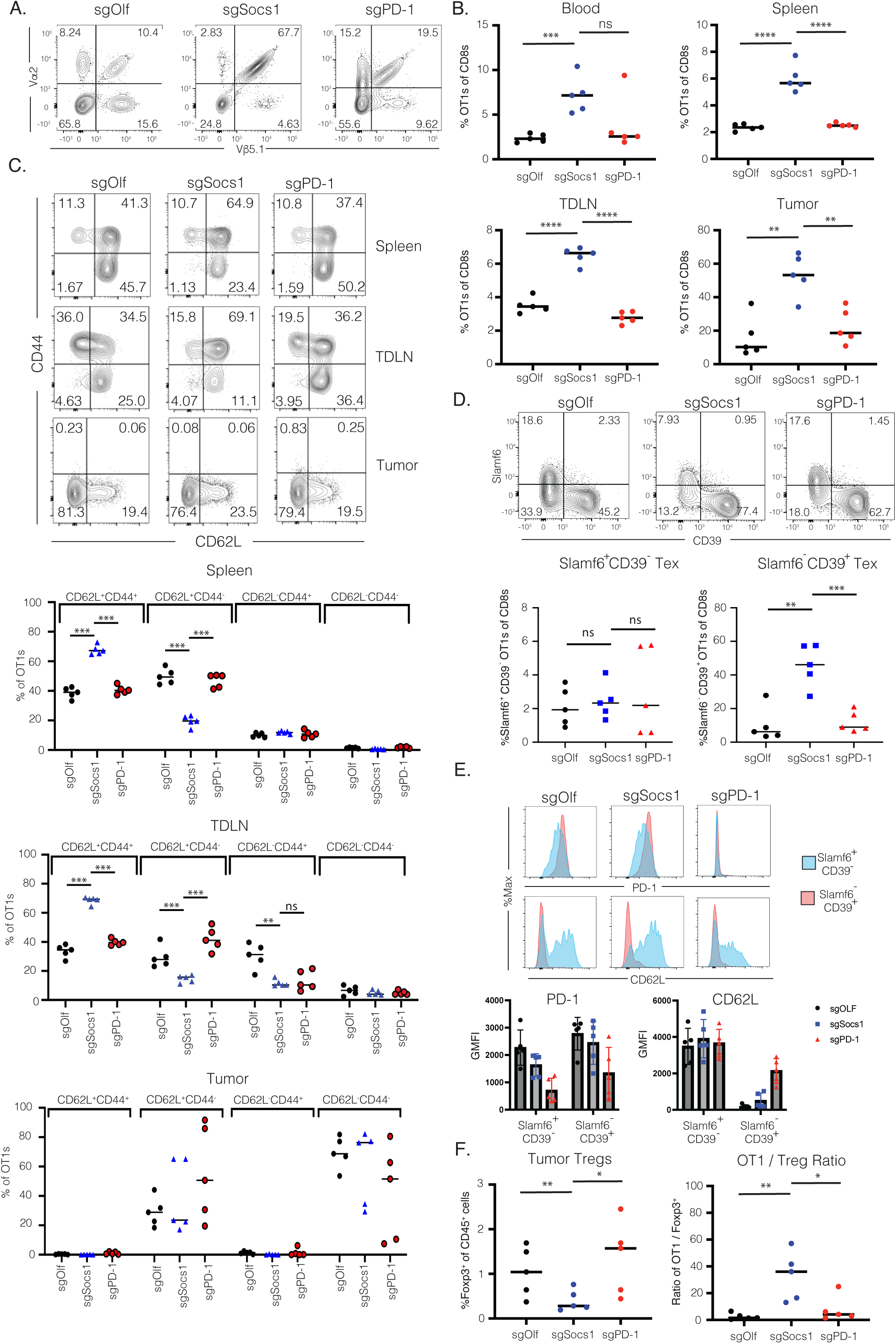
Inactivation of SOCS1 in transferred CD8 T cells enhances their accumulation as CD44^+^CD62L^+^ T_cm_ cells within lymphoid organs and Slamf6^-^CD39^+^PD-1^hi^ Tex cells in tumors while depleting intratumoral Tregs. C57BL/6 mice bearing 100mm^3^ B16-Ova tumor cells bearing a median size of 100mm^3^ were treated with 3x10^6^ SOCS1 (sgSocs1), PD1 (sgPD-1) or OLF1 (sgOlf) engineered OT1s. **(A)** OT1 frequency in tumors 7 days following transfer was determined by quantifying the frequency of CD8^+^Vα2^+^Vβ5.1^+^ cells within the CD8 T cell compartment, with representative mouse shown. **(B)** Frequency of CD8^+^Vα2^+^Vβ5.1^+^ cells in the blood, spleen, tumor-draining lymph nodes (TDLNs), and tumor between treatment groups. **(C)** The frequency of OT1s between treatment groups expressing CD62L and/or CD44 are depicted from TDLN, spleen and tumor. **(D)** The expression pattern of Slamf6 and CD39 by OT1s from tumor depicted. **(E)** The expression of PD-1 or CD62L protein expressed by either Slamf6^+^CD39^-^ or Slamf6^-^CD39^+^ intra-tumoral OT1s from each treatment group is depicted from a representative mouse in the top row, with compiled data from individual mice shown in the bottom row. **(F)** The frequency of CD4^+^Foxp3^+^ cells within the TME was quantified in relation to total CD45^+^ cells (graph on the left) and as a ratio of OT1s : Tregs (right). Each symbol reflects and individual mouse, with *** = p value < 0.001; ** = p value < 0.01, * = p value < 0.05 and ns = no significance by Student’s t test between the indicated comparator groups.

In assessing the impact of SOCS1 and PD-1 inactivation in transferred CD8 T cell on the cellular composition of tumor, we observed depletion of Foxp3^+^ T regulatory cells relative to sgOlf OT1s in mice treated with sgSocs1 OT1s but not sgPD-1 OT1 treated animals. This resulted in intra-tumoral OT1 / Treg ratio of 32-fold with sgSocs1 OT1s in comparison to 2.6-fold with sgOlf OT1s and 8-fold with sgPD-1 OT1s (Figure 3F). Of note, the mean tumor volume of sgOlf, sgPD-1 and sgSocs1-treated groups on the day of tumor harvest were 394mm^3^ SEM +/-35, 210mm^3^ SEM +/-110, and 287mm^3^ SEM +/- 81, respectively, suggesting that the depletion of Tregs in the sgSocs1 treatment group is not driven by stronger efficacy and smaller tumor size. We also assessed the impact of treatment groups on intra-tumoral innate cell population frequencies(39). Populations evaluated include CD45^+^CD11b^+^Ly6C^+^Ly6G^+^ neutrophils; CD45^+^CD11b^+^Ly6C^+^Ly6G^-^MHCII^+^F4/80^-^ monocytic myeloid-derived suppressor cells (mMDSCs); CD45^+^CD11b^+^Ly6C^med^Ly6G^-^MHCII^+^F4/80^+^ tumor associated macrophages (TAMs) including CD11c^-^ TAM1 and CD11c^+^ TAM2 subsets; CD45^+^CD11b^+^Ly6C^-^MHCII^+^ CD11c^+^ classical dendritic ells (cDCs) including CD103^+/-^ cDCs and CD11b^+^NKp46^+^ natural killer (NK) cells (Figure S3A-B). The sgSocs1 treatment group was found to increase the presence of neutrophils and decrease the presence of mMDSCs, CD103^+^ cDCs as well as TAM2 subsets from within the tumor (Figure S3B). These results together with the observed depletion of intratumoral Tregs driven by sgSocs1 OT1s indicate that inactivation of SOCS1 in transferred CD8 T cells remodels tumor by in part reducing the frequency of immunosuppressive cell subsets known to impede T cell anti-tumor function.

### SOCS1 is a key checkpoint in the accumulation of Tex^int^ and Tex^eff^ cells from Tex^prog^ subsets within tumors with mechanisms distinct from PD-1

To understand in greater detail how inactivation of SOCS1 impacts the state of transferred CD8 T cells within the tumor, scRNA-Seq (10x Genomics) was performed on CD45^+^ bead-selected cells isolated from B16-Ova tumors harvested 7 days following the transfer of either sgSocs1, sgPD-1 and sgOlf OT1s. Across all treatment groups, 18,794 independent transcriptomes were captured, with expression signatures revealing the presence of major immune cell types including macrophages, DCs, NK cells, B cells, and T cells (Figure S4A). Differences in overt cell type populations between treatment groups were minimal, with the exception being an enrichment of B cells in the sgPD-1 treated group (Figure S4B). Analysis of the CD3^+^ T cell subpopulation comprising 8123 independent transcriptomes revealed 11 distinct clusters (Figure 4A, with Figure S4C depicted cluster cell counts) as defined by T cell cluster marker genes (Figure S4D). The use of 5’ scRNA-Seq permitted detection of OT1 TCR alpha and beta chain sequences within the annotated CD3^+^ clusters, with 2377 OT1s identified (sgOlf: 674; sgSocs1: 1187; and sgPD-1: 516 OT1s detected per treatment group, respectively) and found to localize within four T cell clusters: 2, 3, 5 and 7 (Figure 4B). Detection of discrete OT1s allowed for pairwise comparison of gene expression signatures between treatment groups, with pseudobulk RNA-Seq analysis used to identify differentially expressed genes (DEGs) both induced (DEG^up^)and depleted (DEG^down^) (Figure 4C). Similar numbers of absolute DEGs were observed between all treatment group comparisons (Figure S4E), with the high number of DEGs in the sgSocs1 vs sgPD-1 comparison indicating distinct mechanistic differences in how inactivation of SOCS1 or PD-1 impact CD8 T cell transcriptional state. In the sgSocs1 vs sgOlf comparison, *Gzmb* transcripts were the highest DEG^up^, with *Ccr7*, *Tcf7* and *Slamf6* identified as the strongest DEG^down^ (Figure 4D). When comparing the sgPD-1 OT1s to either sgSocs1 or sgOlf OT1s, *Tox* transcripts encoding for TOX, a transcription factor and master regulator of T cell exhaustion(40) was amongst the strongest DEG^up^ (Figure 4D). GSEA was performed by projecting DEGs from pairwise treatment group comparisons onto published gene expression signatures described by Miller et al to define progenitor, proliferating, effector-like and terminally exhausted Tex subsets transcriptionally and functionally (41). We found that both sgSocs1 and sgPD-1 OT1s enriched in proliferating, effector-like and terminally exhausted Tex expression signatures while depleted in progenitor signatures when compared to sgOlf OT1s (Figure 4E). Conversely, when comparing sgSocs1 versus sgPD-1 OT1s, sgSocs1 was enriched for proliferating and effector-like and depleted of both progenitor and terminally exhausted Tex signatures (Figure 4E). These results together with the observed induction of *Tox* transcripts by sgPD-1 OT1s highlights a mechanistic distinction between the impact of sgSocs1 and sgPD-1 on the transcriptional state of transferred CD8 T cells.

**Figure 4:**
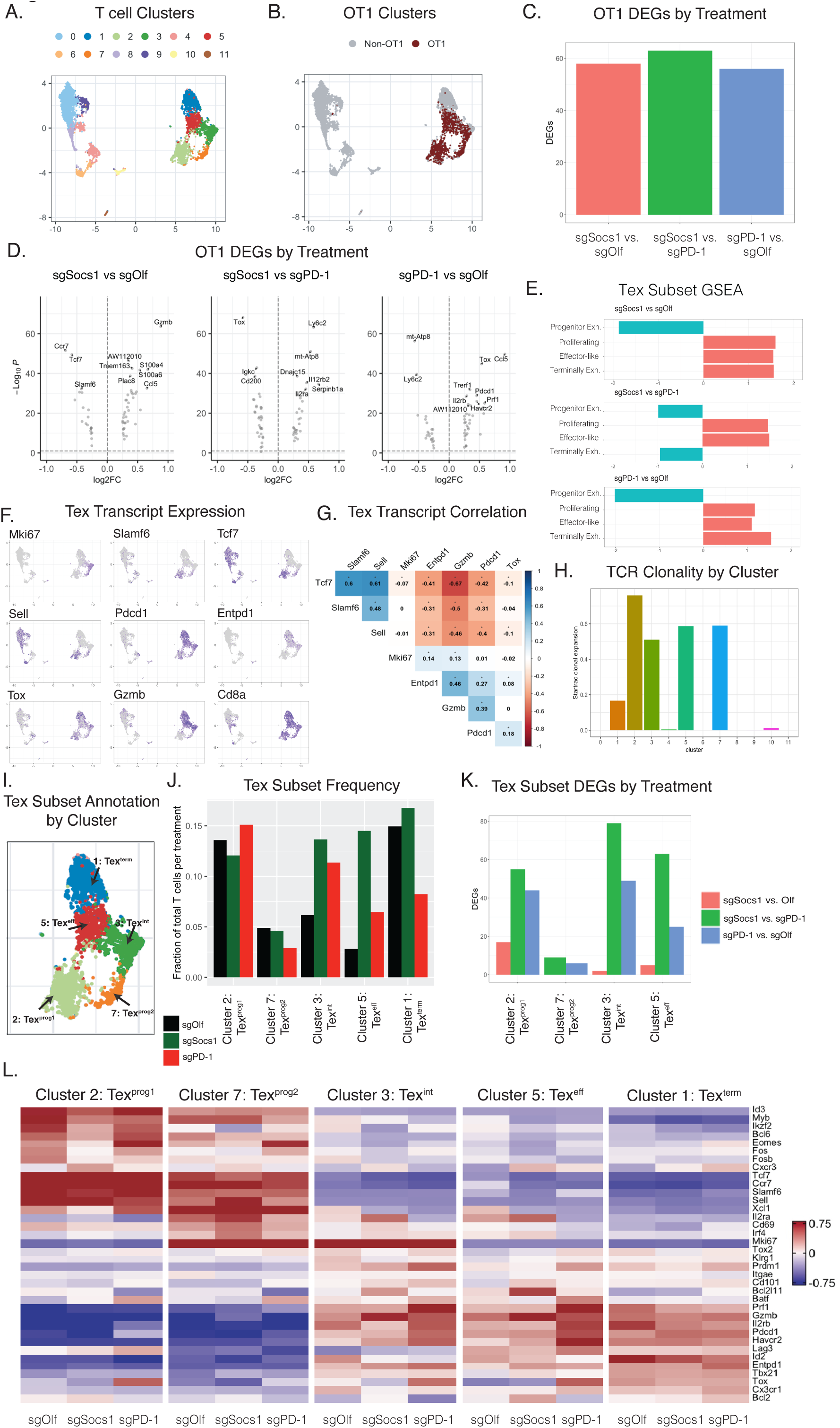
SOCS1 is a key checkpoint in the accumulation of Tex^int^ and Tex^eff^ cells from Tex^prog^ subsets within tumors with mechanisms distinct from PD-1. C57BL/6 mice bearing 100mm^3^ B16-Ova tumor cells bearing a median size of 100mm^3^ were treated with 3x10^6^ SOCS1 (sgSocs1), PD1 (sgPD-1) or OLF1 (sgOlf) engineered OT1s, with editing efficiencies for target genes 82% for sgPD-1, 95% for sgSocs1, and 80% for sgOlf. scRNA-Seq was performed on CD45^+^ cells isolated from the TME from each treatment group. **(A)** UMAP visualization of T cell clusters. **(B)** Projection of OT1s onto T cell clusters based on TCR sequencing. **(C)** Bar plot of treatment DEGs between treatment-group OT1s, adjusted p value < 0.1, abs(avgLog2FC > =0.25). **(D)** Pseudo-bulk analysis on OT1s, with DEGs between treatment groups depicted. **(E)** GSEA by projecting pseudo-bulk DEGs from between indicated treatment groups in (d) onto Miller *et al.* Tex subset gene signatures. **(F)** UMAP visualization depicting the expression of indicated transcripts by T cell clusters **(G)** Correlogram between indicated transcripts (* = p < 0.001) **(H)** STARTRAC TCR clonal expansion by T cell cluster **(I)** Tex subset annotation by cluster. Tex^prog1^ and Tex^prog2^ = progenitor subsets 1 and 2; Tex^int^ = intermediate; T^eff^ = effector-like, and Tex^term^ = terminally differentiated Tex subsets. **(J)** Tex subset frequency by treatment group. Clusters 2, 7, 3 and 5 reflect OT1 frequencies, with Cluster 1 containing non-OT1 cells and included for reference. **(K)** Number of DEGs within each Tex subset as indicated, and between depicted treatment groups **(L)** Heatmap of Tex subset-defining transcripts by subset and by treatment group.

Our FACS analysis of tumor-infiltrating OT1s found that inactivation of SOCS1 in transferred CD8 T cells did not impact Slamf6^+^CD39^-^PD-1^med^ Tex^prog^ cell frequency while enhancing the accumulation of Slamf6^-^CD39^+^PD-1^hi^ cells containing Tex^int^, Tex^eff^ and Tex^term^ cells (Figure 3D). We next sought to query CD8 Tex states encompassed within these T cell clusters in more detail. To ensure that our analysis captured all T cell clusters responding to cognate tumor antigen and including non-OT1 endogenous T cells, we identified T cell clusters with high TCR clonality and thus undergoing clonal expansion using STARTRAC(42). We observed expected strong TCR clonality within the four OT1-enriched clusters and identified cluster 1 as also characterized by high TCR clonality, even with few OT1s present (Figure 4H). We focused our analysis of Tex subsets on these five clusters hereafter. These clusters demonstrated ubiquitous expression of *Cd8a* transcripts with heterogenous expression of Tex-state defining transcripts. Cluster 1 was represented by *Pdcd1, Entpd1, Tox* and *Gzmb* transcripts; cluster 2 represented by *Slamf6, Tcf7,* and *Sell*; cluster 3 represented by *Mki67, Pdcd1* and *Gzmb*; cluster 5 represented by *Pdcd1, Entpd1,* and *Gzmb*; and cluster 7 represented by *Mki67, Slamf6, Tcf7,* and *Sell* (Figure 4F). Strong positive correlation of transcript expression was observed between *Tcf7, Slamf6* and *Sell* and between *Entpd1, Gzmb* and *Pdcd1*, with *Tcf7, Slamf6* and *Sell* in turn all showing strong inverse correlation with *Entpd1, Gzmb* and *Pdcd1* transcripts (Figure 4G). The inverse correlation between *Slamf6* and *Entpd1* transcripts, which encode for Slamf6 and CD39 proteins, respectively, is consistent with our observations in Figure 3D, in which Slamf6^+^CD39^-^ and Slamf6^-^CD39^+^ populations could be clearly demarcated.

To align Tex cell states to each cluster undergoing clonal expansion, we compared cluster expression signatures to published RNA-Seq and scRNA-Seq datasets derived from chronic viral infection and tumor settings and used to define exhausted T cells subsets based on transcriptional and functional profiles (41, 43). Each cluster was projected on to gene signatures previously used to define progenitor Tex cells (Tex^prog1^), proliferating subsets encompassing both progenitor Tex^prog2^ and intermediate Tex^int^ cells, effector-like Tex^eff^ cells, and terminally differentiated Tex^term^ cells (Figure S4F). We were able to discern the continuum of the Tex differentiation pathway within our five identified clusters (Figure 4I,L) (43–45), and identified Tex^prog1^ subset as quiescent *Slamf6^+^Tcf7^+^* cells (cluster 2) and Tex^prog2^ subset as proliferating *Mki67^+^Slamf6^+^Tcf7^+^* cells (cluster 7) with both clusters reflecting TCF-1 expressing stem-like progenitor Tex cells. A population of proliferating Tex^int^ *Mki67^+^Gzmb^+^Entpd1^+^* cells (cluster 3) and quiescent Tex^eff^ *Mki67^-^Gzmb^+^Entpd1^+^* cells (cluster 5) reflecting Tex cells with effector capabilities characterized by expression of *Gzmb* transcripts were also identified, as well as a population of quiescent *Gzmb^+^Entpd1^+^Tox^+^* cells (cluster 1) reflecting terminally differentiated (Tex^term^) Tex cells as characterized by high expression of *Tox* transcripts. Of note, few OT1s were present in the Tex^term^ cluster. Tex^prog1^, Tex^eff^ and Tex^term^ clusters were found to be enriched for TCR activation pathway gene signatures and with Tex^prog2^ and Tex^int^ conversely showing depletion, correlating with quiescent state of the former and proliferating state of the latter Tex subsets (Figure S4G) (46). (47)The impact of SOCS1 or PD-1 inactivation on Tex subsets was next evaluated. SOCS1 or PD-1 inactivation had no impact on overall frequencies of either Tex^prog1^ or Tex^prog2^ subsets reflecting TCF1-expressing progenitor Tex cells (Figure 4J). Given heightened expression of *Slamf6* and lack of expression of *Entpd1* transcripts by Tex^prog1^ and Tex^prog2^ subsets, this is consistent with our observation that treatment did not impact frequencies of Slamf6^+^CD39^-^ OT1s (Figure 3D). However, inactivation of SOCS1 in the sgSocs1 OT1 group drove a statistically significant enrichment of both Tex^int^ (p-value = 7.67x10^-17^) and Tex^eff^ subsets (p-value = 1.7x10^-62^) reflecting proliferating and quiescent Tex subsets with effector capabilities (Figure 4J). As Tex^int^ and Tex^eff^ subsets do not express *Slamf6* yet express *Entpd1* transcripts, this finding is also consistent with our observed increase in frequency of Slamf6^-^CD39^+^ OT1s in the sgSocs1 treatment group. CD4^+^Foxp3^+^ Tregs localized to Cluster 10 (Figure S4H), with this cluster depleted in the sgSocs1 treatment group (p-value = 1.33x10^-05^), consistent with our FACS data (Figure 3F). To determine how treatment groups impacted gene expression signatures within discrete Tex clusters, we examined DEGs between treatments within Tex subsets (Figure 4K) and evaluated the expression patterns of a curated list of transcripts involved in defining Tex state (Figure 4L). Interestingly, pairwise comparison of the sgOlf vs sgSocs1 yielded few DEGs within each Tex cluster (Figure 4K, Figure S4J). For Tex subset DEGs in the sgPD-1 vs sgOlf and sgSocs1 vs sgPD-1 comparison, a far greater overall number of DEGs were identified (Figure 4K), with *Tox* again a top DEG^up^ in the sgPD-1 group across multiple Tex subsets (Figure 4L, Figure S4J). We note that the detection of transcripts aligning to the *Pdcd1* gene in the sgPD-1 group (Figure 4L), despite an editing efficiency of 82% in sgPD-1 OT1s with a sgRNA targeting exon 2 of the *Pdcd1* gene, is likely due to our use of 5’ scRNA-Seq permitting detection of transcribed yet edited transcripts from the *Pdcd1* gene.

These scRNA-Seq data confirm and extend our FACS analysis in demonstrating how inactivating SOCS1 impacts the trajectory of transferred CD8 T cells as Tex subsets within tumors in comparison to PD-1. Tex^int^ and Tex^eff^ cells are known to differentiate from Tex^prog^ subsets (47), with our data suggesting that SOCS1 serves as a key checkpoint in the accumulation of Tex^int^ and Tex^eff^ cells from Tex^prog^ cells. Inactivation of SOCS1 had little impact on the transcriptional states of Tex subsets, with the most pronounced impact of SOCS1 inactivation being the overt increase of Tex^int^ and Tex^eff^ subsets exhibiting high expression of *Gzmb* transcripts. The impact of SOCS1 inactivation on transferred CD8 T cells is mechanistically from PD-1 since while PD-1 also enriched for effector Tex subsets, albeit to a lesser degree then inactivation of SOCS1, inactivation of PD-1 also enriched for a terminally exhausted gene signature including heightened expression of *Tox* transcripts.

### A CRISPR tiling screen in primary human T cells identifies highly potent sgRNAs for therapeutic use targeting the SH2 domain of SOCS1

To extend our observation that inactivation of SOCS1 enhances the anti-tumor function of TIL with the objective of applying these findings for therapeutic use, we next identified potent and selective sgRNAs targeting the human *SOCS1* gene. We systematically evaluated all potential sgRNAs targeting the coding sequence (CDS) of SOCS1 by first eliminating sgRNAs predicted to cut multiple sites in the genome and then performing a CRISPR screen using a sgRNA tiling library that included 134 SOCS1-targeting sgRNAs with unique genome-cutting sites. The CRISPR tiling screen approach can serve to inform the functional impact of targeting various SOCS1 protein domains with candidate sgRNAs(48). Using our observation that SOCS1 restrains accumulation of activated human T cells in the presence of IL-2 in vitro, the sgRNA tiling library was introduced into activated primary human CD3^+^ T cells by lentivirus transduction with Cas9. The sgRNA Lib^+^ T cells were then expanded in the presence of IL-2 (Figure 5A). sgRNAs targeting the KIR, the extended SH2 subdomain (ESS) and the SH2 domain of SOCS1 robustly enriched in comparison to sgRNAs targeting other regions of the protein, including the NTD and SOCS Box domains (Figure 5B). The KIR domain has been found to directly block the substrate-binding groove of JAKs and the SH2 domain mediating binding to JAK proteins, with these domains having a role in SOCS1 function(49). Our data indicated that sgRNAs targeting the KIR, ESS and SH2 domain of SOCS1 provide the most potent inactivation of SOCS1 protein function.

**Figure 5:**
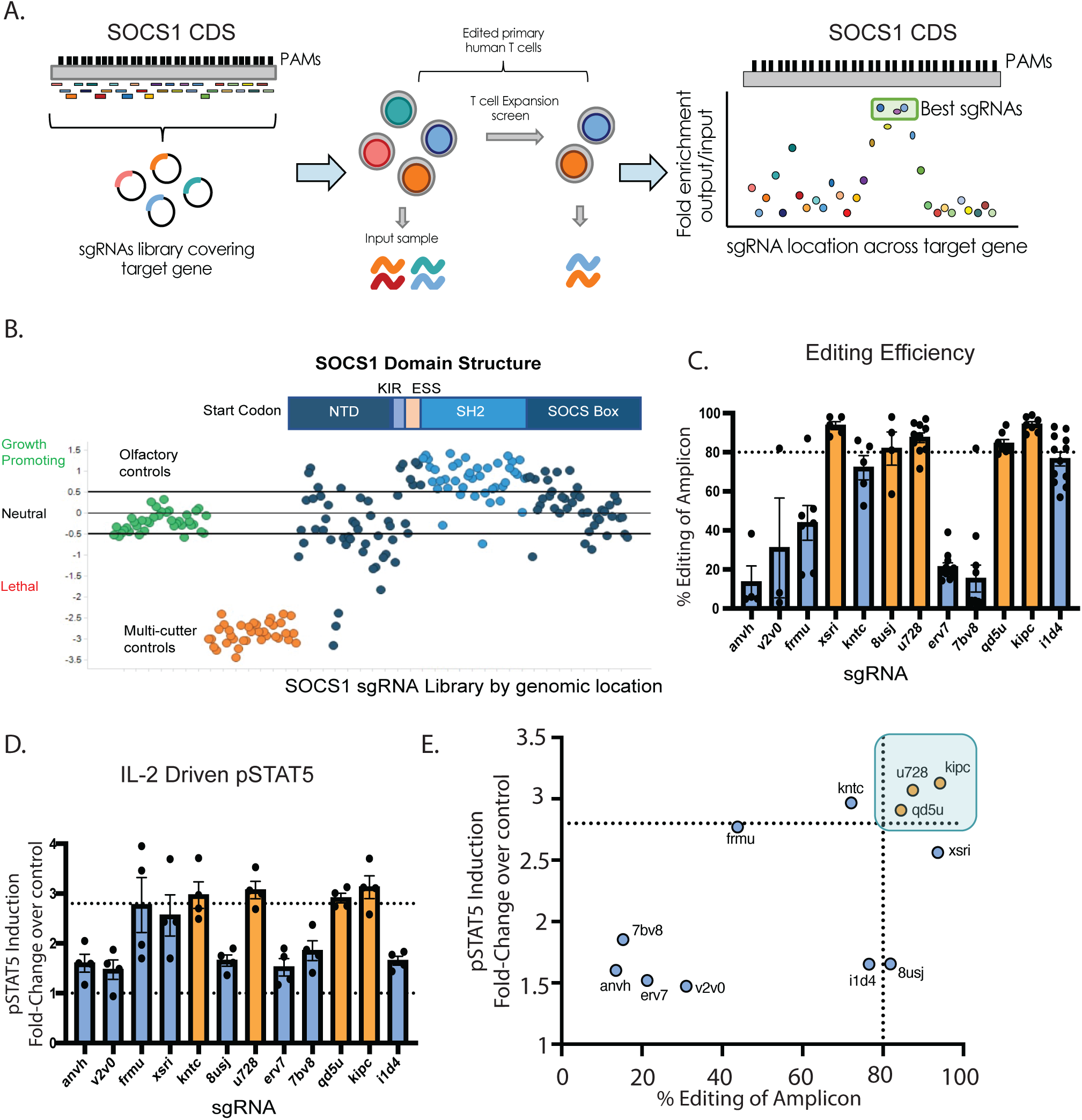
A CRISPR tiling screen in primary human T cells identifies highly potent sgRNAs for therapeutic use targeting the SH2 domain of SOCS1. **(A)** Experimental schematic of the CRISPR tiling screen to discover potent SOCS1 sgRNAs. Following in silico removal of sgRNAs predicted to target multiple sites in the genome, a sgRNA library targeting every possible Cas9 cut site of the SOCS1 coding region (CDS) based on the trinucleotide NGG PAM sequence together with controls was introduced by lentiviral transduction into activated primary human T cells with IL-2. Following introduction of Cas9, sgRNA Lib^+^ T cells were expanded in the presence of IL-2, with the distribution of sgRNAs following expansion evaluated and compared to input. **(B)** SOCS1 CRISPR tiling screen results. sgRNAs targeting the olfactory genes are in green, genome multi-cutters in orange, and sgRNAs targeting the SOCS1 CDS are blue. The SOCS1 protein domain structure is depicted above, with NTD = N-terminal domain, KIR = kinase inhibitory region, ESS= extended SH2 subdomain, and with the SH2 and SOCS Box domains labelled. sgRNAs targeting the SH2 domain of SOCS1 are depicted in light blue. **(C)** The editing efficiency of top sgRNAs identified in (b) was assessed by electroporation of Cas9/sgRNA RNPs into activated primary human T cells in an arrayed format, with editing efficiency of the cut site quantified by Amp-Seq. sgRNAs are labelled along the x-axis, with red bars depicting sgRNAs achieving editing efficiency of 80% or greater. **(D)** Activated human primary T cells were edited with Cas9/sgRNA RNPs targeting either SOCS1 or Olfactory genes in an arrayed format, with edited T cells stimulated with IL-2 and phospho-STAT5 signals quantified by FACS and depicted as fold-change over sgOlf control. sgRNAs are labelled along the x-axis, with red bars depicting SOCS1 sgRNAs achieving the targeted IL-2-mediated increase in pSTAT5. **(E)** A comparison between editing efficiency (x-axis) and pSTAT induction fold-change (y-axis) of evaluated sgRNAs, with the u728, kipc, and qd5u sgRNAs identified as the most potent sgRNAs targeting SOCS1 based on editing efficiency and functional potency.

We next quantified the editing efficiency of the top 12 of the highest ranking sgRNA’s identified in the SOCS1 CRISPR tiling screen by electroporating primary human T cells with sgRNA/Cas9 RNPs in an arrayed format and determining editing efficiency by Amp-Seq. We found that multiple sgRNAs were able to robustly achieve greater than 80% editing efficiency across at least four independent donors (Figure 5C). As SOCS1 is a negative regulator of IL-2 signaling, we also evaluated the functional potency of candidate sgRNAs in a human T cell pSTAT5 assay, using IL-2 as a stimulus. Multiple sgRNAs were able to drive enhanced IL-2-mediated pSTAT5 signals in primary human T cells in comparison to unedited controls (Figure 5D), with editing efficiency strongly correlating with pSTAT5 enhancement (Figure 5E).

To identify potent SOCS1 sgRNAs with minimal off-target edits, the most potent sgRNAs were additionally interrogated for their off-target profile using GUIDE-Seq(50). This method can detect off-target dsODN integration events occurring at frequencies as low as 0.1% relative to on-target insertion reads. Across six independent donors and four repeat studies, we found that the u728 sgRNA targeting the 3’ end of the SH2 domain of SOCS1 displayed strong editing efficiency with 22 potential off-target sites identified (Figure S5A). As GUIDE-Seq is a useful tool to discover genomic cut-sites, we next verified the on- and off-target editing profile of the u728 guide at the native loci in human melanoma-derived TIL using Amp-Seq. In addition to observing over 90% on-target editing efficiency of SOCS1 in TIL, we found that only 1 of the 22 potential off-target sites identified by GUIDE-Seq was found to be statistically different between u728-edited versus unedited TIL by Amp-Seq (Figure S5C). Importantly, this confirmed off-target site, which occurs at position chr7:133126575 in the GRCh38 assembly (Figure S5D), is located within an intergenic region with no known function, and thus does not represent a significant genotoxicity risk. Based on its strong potency and clean off-target profile, we prioritized the use of u728 sgRNA to be used for the inactivation of the SOCS1 gene in TIL for therapeutic use.

### CRISPR/Cas9 inactivation of the *SOCS1* gene in human TIL drives enhanced functionality and pSTAT4 sensitivity to IL-12

We next used the u728 sgRNA to manufacture KSQ-001, an autologous engineered TIL therapy, by inactivating the *SOCS1* gene by electroporation of sgRNA/Cas9 RNPs (Figure 6A). Across 26 independent datasets and 15 donors, we consistently observed editing efficiencies of the *SOCS1* gene of over 80% in KSQ-001 (Figure 6B), with editing of the *SOCS1* gene resulting in complete depletion of SOCS1 protein in KSQ-001 manufactured from both NSCLC and melanoma donors (Figure 6C). We observed that the electroporation-based engineering step did not impact KSQ-001 expansion during the REP (Figure S6A) or viability following cryopreservation and thaw (Figure S6B), or activation-induced cell death (AICD) in comparison to unedited TIL (Figure S6C). KSQ-001 consisted of CD3^+^ cells with no detectable residual tumor cells, with patient-unique frequency of CD4 and CD8 T cells not impacted by inactivation of SOCS1 (Figure S6D). KSQ-001 displayed a CD45RO^+^CCR7^-^ T_em_ phenotype following harvest, similar to TIL (Figure S6E), and demonstrated enhanced expression of CD25 in comparison to TIL (Figure S6F)

**Figure 6:**
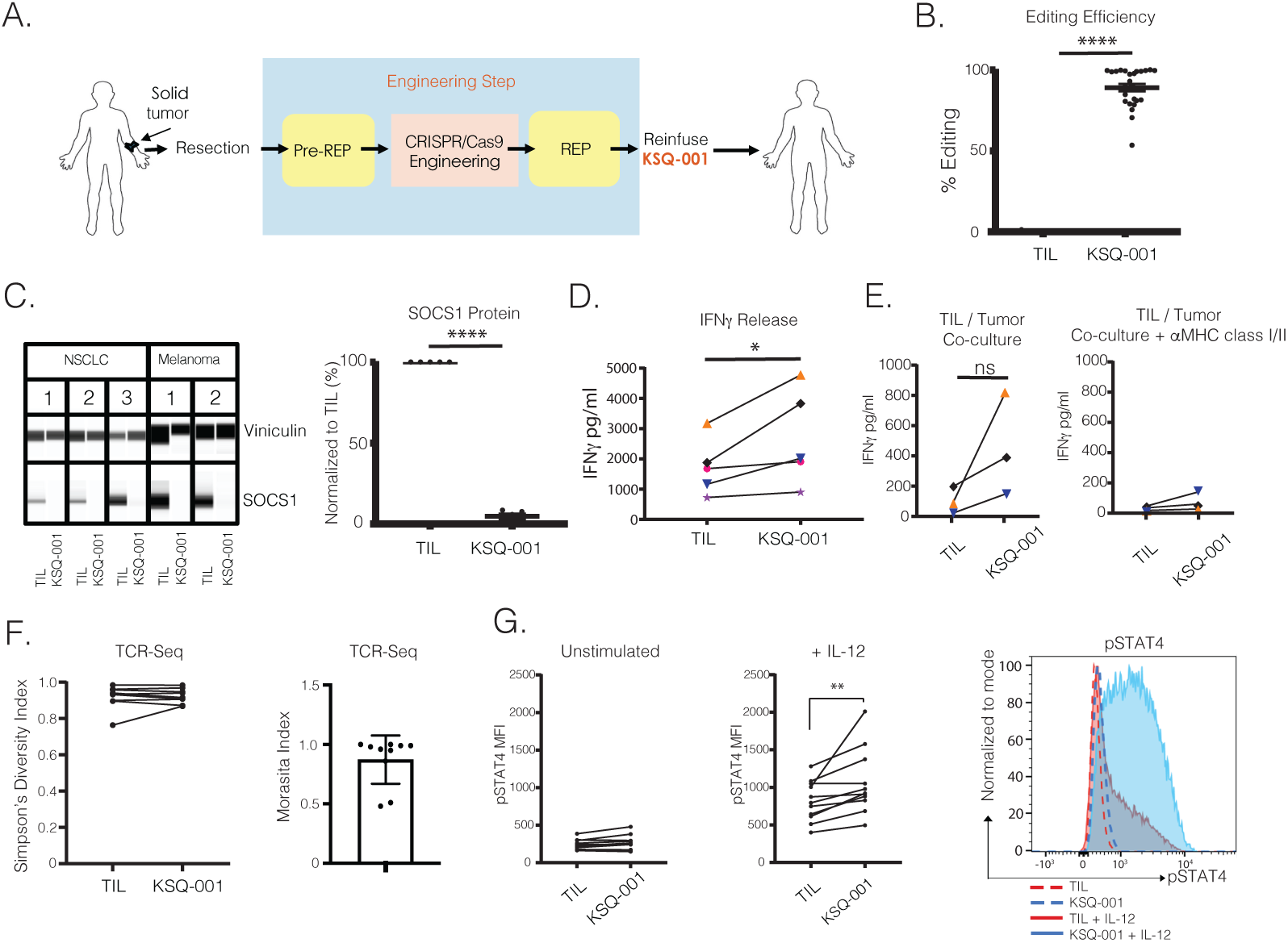
CRISPR/Cas9 inactivation of the *SOCS1* gene in human TIL drives enhanced functionality and pSTAT4 sensitivity to IL-12. (**A**) Schematic of the process to manufacture KSQ-001, a CRISPR/Cas9-engineered TIL (eTIL) containing inactivation of SOCS1. Solid tumors are surgically resected from patients, with TIL extracted from fragmented tumor by culturing in the presence of IL-2 in a pre-REP. Following the pre-REP, u728 / Cas9 protein RNPs are introduced into the TIL by electroporation, followed by expansion in a REP in the presence of irradiated PBMC feeders, IL-2 and OKT3. KSQ-001 is then harvested and cryopreserved for transport to a patient for infusion (**B**) Sequencing of PCR amplicons from the u728 cut site is displayed from 15 independent donors and 26 independent datasets. (**C**) Wes-based detection of SOCS1 protein in TIL and KSQ-001 samples from the indicated tumor type. SOCS1 protein was detected as a single band at ∼33kd. Vinculin (117kd) was used as loading control. (**D**) IFNγ release by donor-paired TIL and KSQ-001 following cryopreservation and thaw, and activation with anti-CD3 tetramers. (**E**) TIL reactivity to autologous tumor digests was assessed by co-culture, with IFNγ production quantified in the absence (left) or presence of anti-MHC class I and class II blocking antibodies. (**F**) The diversity and overlap of donor-paired TIL and KSQ-001 CDR3 TCR repertoire results as quantified by FR3AK-U-Seq are displayed as Simpson’s Diversity Index and Morisita Index, respectively. (**G**) TIL or KSQ-001 were stimulated by IL-12 for 1hr (middle) in comparison to an unstimulated control (left). pSTAT4 MFI gated on donor-paired CD3^+^ TIL and KSQ-001 is displayed. A representative pSTAT4 FACS plot from a single donor is depicted (right). For Figure 6B-G, each dot represents individual donor. Statistical analysis were performed using a Students’ *t* test, with ns = no significance, * = p value < 0.05, ** = p value < 0.01; *** = p value < 0.001, and **** = p value < 0.0001.

IFNγ release by TIL upon activation through the TCR is used as a surrogate measure of potency. We observed that KSQ-001 produced higher levels of IFNγ upon TCR activation in comparison to TIL (Figure 6D). To confirm that the enhancement of IFNγ production was driven by inactivation of SOCS1 and not an artifact of electroporation, we also observed enhanced IFNγ production in KSQ-001 in comparison to TIL in which the *Olf1A* gene had been inactivated (Figure S6G). We next evaluated how KSQ-001 responded to co-culture with autologous tumor digests, with KSQ-001 maintaining ability to produce IFNγ in comparison to TIL, with cytokine release dependent upon MHC class I and II as inclusion of pooled antagonistic antibodies inhibited all IFNγ production (Figure 6E). These findings demonstrate that KSQ-001 retains specificity for autologous tumor. TIL and KSQ-001 retained similar polyfunctionality for the production of IL-2, IFNγ and TNFα cytokines, with no differences observed in the impact of SOCS1 inactivation in KSQ-001 across both CD8 and CD4 T subsets across multiple donors (Figure S6H). The TCR polyclonality of TIL is an important feature thought to drive clinical efficacy by providing comprehensive coverage of all tumor antigens specificities. We therefore wanted to assess whether SOCS1 editing impacted the polyclonality of KSQ-001. We found that KSQ-001 maintained high TCR diversity based on Simpson’s Diversity Index and showed a high degree of TCR overlap with TIL derived from the same donor based on the Morisita Index values (Figure 6F), thus confirming that inactivation of *SOCS1* has no overt impact on the overall TCR repertoire of KSQ-001.

Given the role of SOCS1 as a negative regulator of cytokine signals, we evaluated the sensitivity of KSQ-001 to a panel of cytokines including IL-2, IL-12, and IL-15 by quantifying the levels of pSTAT5 signals for IL-2 and IL-15, and pSTAT4 for IL-12. Despite inactivation of SOCS1 by the u728 sgRNA strongly enhancing pSTAT5 signals in primary human T cells in response to IL-2 (Figure 5D), KSQ-001 surprisingly did not display enhanced pSTAT5 signals in comparison to TIL in response to IL-2 (Figure S7A) or IL-15 (Figure S7B) when profiled directly from REP. However, KSQ-001 did display heightened responsiveness to IL-12 through increased pSTAT4 accumulation (Figure 6G), suggestive of SOCS1-dependent pharmacology. We reasoned the disconnect between IL-2/pSTAT5 hypersensitivity observed with SOCS1-edited primary T cells but not KSQ-001 may be driven by the long time duration KSQ-001 spent in the presence of IL-2 and TCR activation during manufacture (∼27 days), which may induce compensatory mechanisms that blunt SOCS1-driven pSTAT5 signals in response to IL-2 and IL-15, while sparing pSTAT4 responses to IL-12. In support of this hypothesis, we found that *SOCS1* edited human CD3^+^ T cells activated with anti-CD3/CD28/CD2 tetramers and IL-2 displayed enhanced pSTAT5 signals 7 days post-activation (Figure S7C) yet failed to display pSTAT5 hypersensitivity to IL-2 after 13 days of activation (Figure S7D). These data also demonstrate that the lack of IL-2/pSTAT5 hypersensitivity observed with KSQ-001 is not driven by a fundamental difference between the biology of PBMC-derived T cells and TIL.

### Enhanced IL-2 dependent engraftment and anti-tumor activity by *SOCS1*-edited TIL following transfer into a solid tumor model

Based on the findings above, we hypothesized that once KSQ-001 was removed from an environment replete with IL-2 and TCR activation, such as following infusion into a host, the compensatory mechanisms at play might reverse and KSQ-001 could re-gain hypersensitivity to IL-2. To test this hypothesis, we set out to evaluate the cytokine dependent engraftment of KSQ-001 and TIL following adoptive transfer into immunodeficient mouse models. KSQ-001 and TIL were adoptively transferred into NOG or human IL-2 transgenic NOG mice (hIL-2 Tg NOG), with the engraftment of TIL evaluated over time. We found that both KSQ-001 and TIL failed to engraft in NOG mice, indicating there is a baseline level of cytokine support needed to support TIL and KSQ-001 engraftment not present in NOG mice. In the hIL-2 Tg NOG mouse strain, KSQ-001 displayed a marked enhancement in engraftment when compared to TIL, with KSQ-001 comprising 95% of total viable blood cells in comparison to 3% for TIL, suggesting that inactivation of the *SOCS1* gene does indeed drive enhanced IL-2 signals in KSQ-001 when evaluated in an in vivo setting. Expansion of KSQ-001 in hIL-2 Tg NOG mice occured between days 7-14 following transfer, with KSQ-001 presenting as primarily CD8 T cells (Figure 7A). We observed donor heterogeneity in CD4 versus CD8 bias following engraftment in this experimental setting. In a clinical setting, high dose (HD) IL-2 is administered i.v. following TIL infusion, with TIL sustained by endogenous cytokine sinks following cessation of HD IL-2 administration. As the hIL-2 Tg NOG mouse model reflects chronic exposure of TIL to HD hIL-2, we sought to evaluate the ability of KSQ-001 to engraft in two additional settings that may better reflect the clinical setting. We first evaluated engraftment and persistence following daily i.p. administration of 45,000U IL-2 over 14 days, with KSQ-001 again observed to engraft and persist to a greater degree than TIL (Figure 7b). Second, we sought to model the clinical setting where lymphodepleted patients are administered up to three days of high-dose IL-2 following TIL infusion whereupon TIL then rely upon endogenous cytokine sinks, including IL-15, for persistence. We assessed TIL and KSQ-001 engraftment following transfer into human IL-15-Tg NOG over 14 days, with 45,000U of exogenous IL-2 administered over the first three days, with NSG mice included as controls. KSQ-001 again engrafted and accumulated to a greater degree than TIL in this setting, with both KSQ-001 and TIL showing marked improvement in engraftment and persistence in the hIL-15-Tg models in comparison NSG (Figure 7c). These data collectively demonstrate that KSQ-001 possesses hyperresponsiveness to cytokine signals in comparison to TIL following transfer in settings designed to model the clinical treatment of TIL.

**Figure 7:**
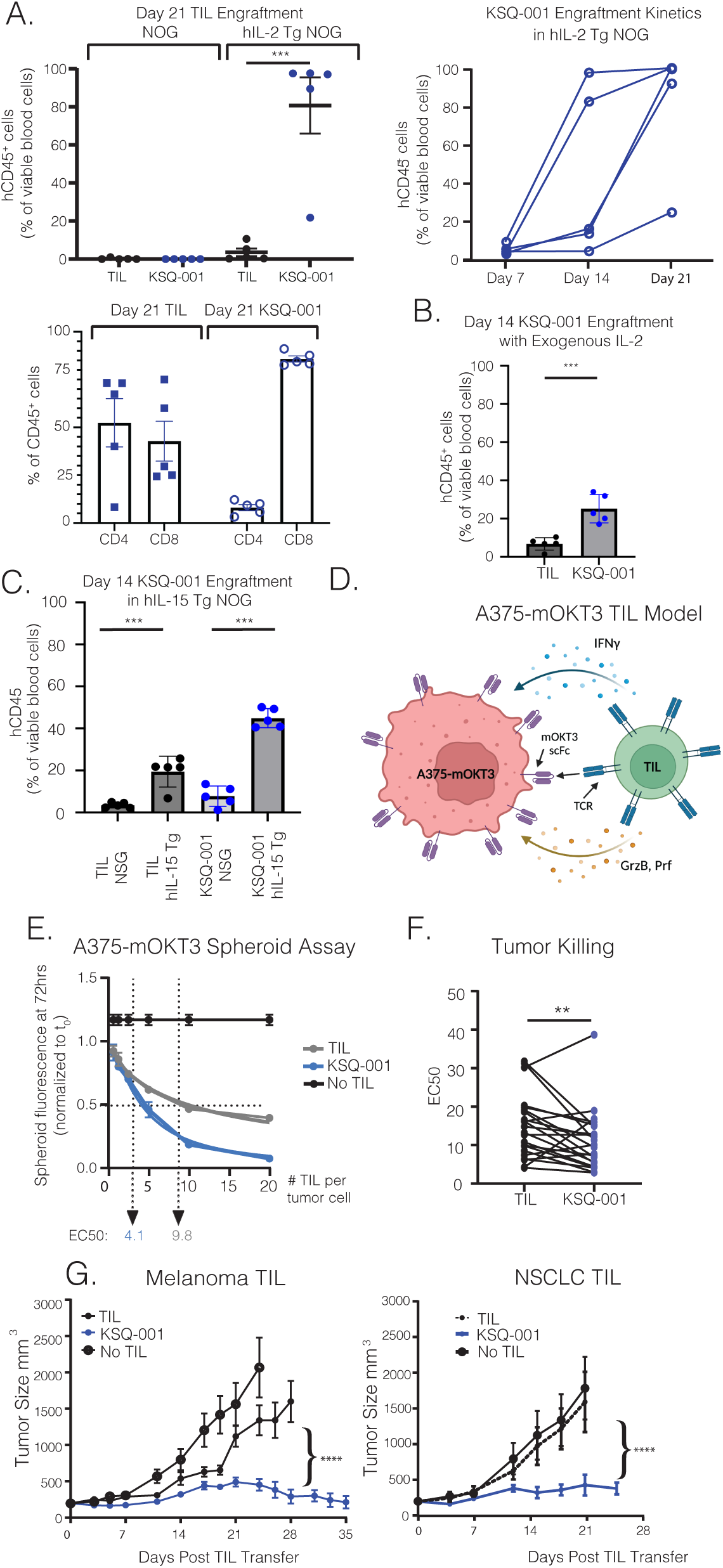
Enhanced IL-2 dependent engraftment and anti-tumor activity by *SOCS1*-edited TIL following transfer into a solid tumor model. **(A)** TIL or KSQ-001 were adoptively transferred into NOG or hIL-2 Tg NOG mice, with engraftment evaluated over time by quantifying the frequency of human CD45^+^ cells present within the peripheral blood. Top row, left: frequency of human CD45+ cells by mouse strain and treatment group on Day 21. Top row, right: frequency of KSQ-001 in hIL-2 Tg NOG mice over time. Bottom row: frequency of CD4 and CD8 T cells from TIL or KSQ-001 on Day 21 in hIL-2 Tg mice. **(B)** TIL or KSQ-001 were adoptively transferred into NOG with 45,000U of human IL-2 administered i.p. daily, with engraftment evaluated at Day 14. **(C)** TIL or KSQ-001 were adoptively transferred into hIL-15 Tg NOG mice with 45,000U of human IL-2 administered i.p. daily for three days, with engraftment evaluated at Day 14, and with NSG mice used as a comparator (**D**) Schematic of the mOKT3-A375 / TIL model. A375 melanoma cells were engineered express either high or low affinity membrane associated OKT3 scFv binding domains which bind and agonize CD3 expressed by TIL. TIL generate an anti-tumor cytolytic response through production of IFNγ and release of cytolytic granules. (**E**) TIL were co-cultured with low-affinity A375-mOKT3 spheroids at various effector to target (E:T) ratios, with tumor killing assessed by Incucyte assay over time. EC50 kill curve at 72 hours from a representative donor is shown. **(F)** EC50 values from 17 donors and 22 independent paired TIL and KSQ-001 samples is shown, with each dot representing an individual paired donor. Statistical analysis was done using two-tailed paired T test. **(G)** hIL-2 Tg NOG mice bearing high-affinity A375-mOKT3 tumors ∼100mm^3^ in size were treated with donor-paired sgOlf-edited melanoma TIL or unengineered NSCLC TIL versus donor-paired KSQ-001, with tumor growth assessed over the indicated time-points. A Student’s *t* test was used to evaluate statistical significance between treatment groups in Figure 7A,B, C and G, and a 2-way ANOVA was used in Figure 7G between sgOlf TIL or TIL vs KSQ-001, with ** = p value < 0.01, *** = p value < 0.001, and **** = p value < 0.0001.

A particular challenge with evaluating the anti-tumor activity of TIL in in vivo animal models is the patient-to-patient uniqueness and high polyclonality possessed by the TCR repertoire of TIL. We developed an MHC-independent tumor xenograft model wherein TIL manufactured across donors would recognize and generate a measurable anti-tumor response. We engineered the A375 human melanoma cell line with membrane-bound OKT3 (mOKT3) of both high (mOKT3) and low (mOKT3^LT^) affinity. Polyclonal human T cells and TIL, irrespective of donor, can engage mOKT3, become activated and thus generate a cytolytic anti-tumor immune response (Figure 7D). Given the high avidity of mOKT3 binding domains for the TCR, we used OKT3 engineered to possess lower functional avidity for TCR (OKT^LT^) to better recapitulate the avidity by which TCRs engage pMHC and increase assay sensitivity(51). We first developed an in vitro 3D mOKT3^LT^-A375 tumor spheroid model and performed a dose titration of KSQ-001 or TIL co-cultured with mOKT^LT^-A375 spheroids and observed that KSQ-001 displayed enhanced potency in this assay in comparison to TIL (Figure 7E). This increase in KSQ-001 potency as indicated by EC50 values in comparison to TIL was observed in 19/22 independently evaluated donors (Figure 7F). We next used mOKT3-A375 cells in an in vivo efficacy model in the hIL-2 Tg NOG strain of mice. KSQ-001 displayed markedly enhanced anti-tumor activity compared to either sgOlf-edited TIL or unedited TIL from both melanoma and NSCLC donors (Figure 7G). Collectively, our data demonstrate that CRISPR/Cas9 edited of the *SOCS1* gene imparts a heightened ability of KSQ-001 to engraft and accumulate following transfer in a cytokine-dependent manner, and strongly enhances the in vitro and in vivo activity in a novel, MHC-independent solid tumor model.

## Discussion

To rationally design an functionally-enhanced T cell therapy with improved clinical activity against solid tumors, CRISPR/Cas9 was used to identify SOCS1 and screen for potent and selective SOCS1-targeting sgRNAs for therapeutic use as well as to inactivate SOCS1 in KSQ-001, an engineered TIL (eTIL®) therapy. SOCS1 emerged as a top hit enhancing CD8 T cell enrichment in tumors to a degree greater than other targets known to serve as brakes on T cell function, including CBLB, MAP4K1, NR4A family members and PD-1. SOCS1 has previously been reported to constrain in vivo CD8 T cell anti-tumor function (21, 52–54), in vitro human T cell proliferation (23), and in vivo CD4 T cell expansion (54). Our results confirm and extend these prior studies in multiple ways. First, we demonstrate that inactivation of SOCS1 in T cell therapies can improve anti-tumor activity in animal models refractory to PD-1 inactivation. Second, we find that inactivation of SOCS1 markedly enhances the engraftment and accumulation of transferred CD8 T cells as CD44^+^CD62L^+^ T_cm_ cells within the peripheral lymphoid organs, with SOCS1-edited T cells durably persisting. Importantly, we report that SOCS1 serves as a key checkpoint in the differentiation of intratumoral CD8 T cells from Tex^prog^ to Tex^int^ and Tex^eff^ cells, with strong accumulation of Tex^int^ and Tex^eff^ cell subsets observed, and with inactivation of SOCS1 mechanistically differing from PD-1 on Tex cell state. These results further our understanding of the key signals impacting the differentiation of Tex cells, as discussed below. Lastly, in applying CRISPR/Cas9 to discover a cell therapy candidate for clinical use, we found that sgRNAs targeting the SH2 domain of SOCS1 are the most functionally potent for therapeutic use, and that inactivation of SOCS1 in human TIL leads to enhanced hypersensitivity to IL-12 and hyperresponsiveness in vivo to IL-2 as well as increased anti-tumor effector function.

A recent report by Galy et al. conducted genome-wide CRISPR screens to identify SOCS1 as a key checkpoint of CD4 T cell accumulation and differentiation into polyfunctional Th1 cells including boosting the anti-tumor function of human CAR-T cells containing a mixed population of CD4 and CD8s (54). We report concordance between Galy et al’s description of SOCS1-edited CD8 T cells with our own observations, as both studies observed enhanced anti-tumor activity and accumulation as well as enhanced expression of Granzyme B protein or *Gzmb* transcripts by SOCS1-endited OT1s. As our work focused on the discovery and furthering of the mechanistic underpinnings of SOCS1 in CD8 T cells, our report in the context of Galy et al provides a more comprehensive understanding of how inactivation of SOCS1 may collectively enhance the anti-tumor activity of a drug product containing a heterogeneous mixture of CD4 and CD8 T cells such as TIL or CAR-Ts (54). Given that SOCS1-inactivation enhanced the cytokine polyfunctionality of CD4 T cells with tumors, and as we identified SOCS1 as controlling the transition of Tex^prog^ to Tex^int^ and Tex^eff^ cells in the absence of provided SOCS1-inactivated CD4 T cell help, we predict that the addition of SOCS1-inactivated CD4s and the enhanced cytokines produced within tumor will serve to further augment the impact of SOCS1 inactivation on CD8 Tex state and, by extension, anti-tumor activity. While we did not observe SOCS1 as impacting polyfunctionality of either CD4 or CD8 T cells present within KSQ-001 drug product in comparison to TIL (Figure S6G), this difference likely due to the very different experimental contexts in which polyfunctionality was evaluated within vivo tumor models in comparison to ex vivo expanded human TIL.

A key finding in this study is our observation that SOCS1 serves as a key checkpoint in the transition of Slamf^+^CD39^-^PD-1^med^ Tex^prog^ cells to Slamf6^-^CD39^+^PD-1^hi^ Tex^int^ and Tex^eff^ cells. While not overtly changing the transcriptional state of Tex subsets as was observed with inactivation of PD-1, the role of SOCS1 as a key brake of cytokine signals implicates cytokines as important mediators of the Tex^prog^ to Tex^int^ and Tex^eff^ transition. The role of SOCS1 at this Tex differentiation interface may be a crucial mechanism to leverage, together with enhancement of engraftment and formation of T_cm_ cells in the peripheral lymphoid organs, towards broadly enhancing the anti-tumor activity of T cell therapies, including KSQ-001, a SOCS1-edited TIL therapy. In benchmarking our studies of SOCS1 in syngeneic T cell therapy models to PD-1, we found that while inactivation of PD-1 in transferred CD8 T cells also led to increased Tex^int^ and Tex^eff^ cells, albeit to a lesser degree then SOCS1, a mechanistically distinct transcriptional state emerged that was marked by heightened expression of *Tox* and enrichment of an terminally exhausted gene signature in comparison to SOCS1 inactivation. PD-1 is known to stabilize the Tex^prog^ pool and repress the formation of Tex^term^ subsets, with inactivation leading to erosion of the Tex^prog^ population over time (36). Our data supports that genetic inactivation of PD-1 in tumor-targeting T cells, while enhancing acute anti-tumor function, also accelerates the terminal differentiation of Tex cells. Corroborating this finding include recent reports in a murine T cell chronic exhaustion virus model as well as a clinical study evaluating the persistence of a CRISPR-engineered TCR-Tg T cell targeting NY-ESO1 demonstrated that genetic inactivation of PD-1 in transferred T cells were deleterious for the long-term persistence of cells in the host following transfer (55, 56). This contrasts with SOCS1, where inactivation has little impact on the Tex^prog^ transcriptional signature, and with no evidence of *Tox* transcript induction observed.

We described herein KSQ-001, a CRISPR/Cas-engineered TIL with inactivation of the SOCS1 gene, with discovery facilitated using CRISPR/Cas9 tiling screens to identify sgRNAs for therapeutic use. KSQ-001 could be manufactured with robust inactivation of SOCS1 across multiple donors and solid tumor types, and resembled TIL with respect to cellularity, viability and TCR specificity for tumor. Notably, KSQ-001 displayed markedly enhanced anti-tumor activity in an MHC-independent mOKT3-A375 in vitro spheroid and in vivo tumor model when compared to TIL. TIL is administered to patients following a course of cyclophosphamide and fludarabine-mediated non-myeloablative lymphodepletion (NMA-LD), with patients treated with HD IL-2 immediately following infusion of TIL. NMA-LD is thought to improve the engraftment of TIL by, in part, reducing the number of competing lymphocytes present within the patient and elevating the circulating levels of available cytokines. The role of SOCS1 as an inhibitor of cytokine signals led us to explore KSQ-001 sensitivity to cytokines. Surprisingly, pSTAT5 signals to IL-2 and IL-15 by KSQ-001 evaluated directly from a REP remained similar to TIL. However, and relevant for how KSQ-001 may engraft in patients, KSQ-001 displayed a marked enhancement in IL-2 and IL-2 + IL-15 cytokine-dependent engraftment and expansion in comparison to TIL following transfer into murine hosts. As the inactivation of SOCS1 in KSQ-001 achieves a mechanistically similar goal sought using NMA-LD and HD IL-2 in the clinic, and given the toxicities driven by NMA-LD and IL-2, KSQ-001 may enable exploring the lowering or elimination of NMA-LD and IL-2 in a clinical setting. Supporting this hypothesis is the strong engraftment and long-term persistence of transferred SOCS1-edited T cells in our syngeneic studies conducted in the absence of lymphodepletion or exogenous IL-2. Consistent with our identification of cytokine signaling pathways as playing a dominant role in the function of TIL through SOCS1, Cytokine Inducible SH2 Containing Protein (CISH), which is a close family member of SOCS1, has also recently been described to enhance ACT function (57).

In this study, we describe a systematic approach to using CRISPR/Cas9 in developing an engineered TIL product, KSQ-001, that displays markedly increased anti-tumor activity. Furthermore, we advance our understanding of CD8 T cell biology in the context of T cell differentiation and exhaustion and implicate SOCS1 as a critical checkpoint towards the enhancement of T cell therapies for clinical use.

## Methods

### Methods

#### Mice

Six to nine week old female C57BL/6J, OT1 (C57BL/6-Tg(TcraTcrb)1100Mjb/J) and PMEL (B6.Cg-Thy1^a^/Cy Tg(TcraTcrb)8Rest/J) mice were purchased from The Jackson Laboratory. h-IL2 NOG (NOD.Cg-*Prkdc^scid^* IL-2rg^tm1Sug^Tg(CMV-IL-2)4-2Jic/JicTac) mice were obtained from Taconic Biosciences. To generate Cas9-Tg x TCR-Tg strains for in vivo CRISPR screens, female Cas9 mice were purchased and crossed with OT1 or PMEL mice at The Jackson Laboratory. Mice were allowed to acclimatize for one week and were injected with tumors between 8-12 weeks of age. Mice were maintained in a pathogen-free facility at the Charles River Accelerator and Development Lab (CRADL), Cambridge, MA.

#### Cell Lines

For mouse syngeneic tumor models, the B16-Ova cell line kindly provided by Dr. Randolph Noelle (Dartmouth Medical School, Hanover, NH), and the MC38-gp100 colon cell line kindly provided by Dr. Patrick Hwu (H. Lee Moffitt Cancer Center, Tampa, FL). The A375 human melanoma cell line was obtained from ATCC and engineered to express either a low-affinity or high-affinity membrane associated anti-CD3 binding domain from clone OKT3 (mOKT3) together with RFP. See Supplementary Methods for more information on A375-mOKT3 cell lines.

#### Cell Culture

B16-Ova, MC38-gp100 and A375-mOKT3 cells were cultured in DMEM (Gibco, Cat# 11885076) supplemented with 10% FBS (Gibco, Cat# 10082147) at 37°C in a 5% CO_2_ atmosphere. Cells were passaged two to three times per week to ensure confluency in the flask never surpassed 80%. Tumor cells were harvested in serum-free DMEM during exponential growth phase immediately prior to inoculation.

##### Generation of sgOlf, sgPD-1 and sgSocs1 OT1s and PMELs

Spleens were harvested from OT1 or PMEL mice, placed in 5mL StemCell Buffer and dissociated in GentleMACS C-tubes using a GentleMACS octo dissociator. Following filtration and rinsing of cells, CD8 T cell were isolated using the EasySep^TM^ Mouse CD8+ T Cell Isolation Kit (Stemcell, Cat# 19853) according to the manufacturer’s instructions. Purified CD8 T cells were activated in the presence of 4ng/ml mouse rIL-2 with mouse CD3/CD28 Dynabeads. 48 hours later, Dynabeads were removed, sgRNA / Cas9 RNPs prepared with sgRNA at 22μM and Cas9 at 15μM in IDTE, Buffer T and electroporation enhancer, and electroporated at 1700V, 20ms, 1pulse using a Neon transfection system (LifeTechnologies). Cells were expanded in the presence of 32ng/ml mouse rIL-2 for an additional 48 hours, harvested, and either transferred directly into recipient mice or cryopreserved in 90% FBS + 10% DMSO.

##### Manufacture of TIL

Dissociated Tumor Cells (DTCs) from Discovery Life Sciences were thawed and seeded into 24W Grex (Wilson Wolf) at a density of 1.5x10^6^ to 2x10^6^ /ml in Pre-REP media (RPMI 1640 (Gibco, Cat#11875) supplemented with 10% Human AB Serum (Valley Biomedical, Cat# HP1022HI), 1X Penicillin/Streptomycin Solution (Gibco, Cat#15140-122), 5µg/mL Gentamicin (Gibco, Cat#15710064), 10mM HEPES (Gibco, Cat#15630080), 1X GlutaMax (Gibco, Cat#35050061), 1mM Sodium Pyruvate (Gibco, Cat#11360070) and 3000IU/mL recombinant human IL2 (PeproTech, Cat# 200-02) and cultured at 37^◦^C with 5% CO_2_ for 15-23 days. Every 2-4 days, cells were counted, with a 50% media exchange with 3000IU/mL of IL2 and split. When cell numbers reached 40x10e^6^ pre-REP TIL, TIL were harvested for CRISPR/Cas9 engineering. Briefly, expanded TILs were counted, centrifuged at 300g for 7 minutes and resuspended with MaxCyte electroporation buffer (HyClone Cat#EPB1) according to the manufacturer protocol. Ribonucleoprotein (RNP) master mixes containing Cas9 protein (Aldevron, Cat#9212) and u728 sgRNA or a7mm OLF sgRNA were added to the cell suspension and transferred to processing assembly (MaxCyte), with the specific process assembly selected based on cell numbers. Cells were electroporated on a MaxCyte ExPERT electroporator using the “Optimization #9” program. TIL were then transferred to a recovery plate and each chamber washed with REP media (1:1 ratio of RPMI 1640 and AIMV (Gibco, Cat#12055) supplemented with 5% Human AB Serum and 6000IU/mL recombinant human IL2) twice, with each wash transferred to the recovery plate, which was then incubated at 37 °C for 20 minutes. Following electroporation and recovery, TIL is transferred into G-rex vessel (Wilson Wolf) containing REP media. To initiate the REP, allogenic irradiated PBMC from 5 donors mixed at a 1:1:1:1:1 ratio were mixed with TIL at a 100 to 1 iPBMC : TIL ratio in REP media, with 30ng/mL anti-human CD3 clone OKT3 (Biolegend, Cat# 317347) added. Between Day 1 through Day 14 of REP, a 50% media exchange and 6000IU/mL of IL2, cell count or split was performed based on media change every 2-4 days. On Day 14, cells were harvested and cryopreserved using 100% CryoStor10 (STEMCELL, Cat#07930).

##### IFNγ release assay

100,000 cells per well of KSQ-001 or TIL were plated into a 96well round bottom plate (Falcon, Cat#353077) in REP media, stimulated with a dose response of anti-CD3 tetramer (1µL/mL∼100µL/mL) (STEMCELL, Cat#103109) for 16hr∼24hr, with supernatant IFNγ level measured by enzyme-linked immunosorbent assay (ELISA) following manufacturer’s protocol (Biolegend, Cat# 430104).

##### AICD assay

To evaluate the ability of TIL to undergo activation-induced cell death, KSQ-001 or TIL were stimulated with a dose response of anti-CD3 tetramer (1µL/mL to 100µL/mL) (STEMCELL, Cat#103109). Following activation, cells were harvested and centrifuged at 300G for 3min, stained for CD45 (BD Bioscience, Cat# 563204), CD4 (BD Bioscience, Cat# 560836) and CD8 (BD Bioscience, Cat# 563795), fixed, permeabilized then stained for Caspase3 (Bioscience, Cat# 550914).

##### TIL / Tumor digest co-culture assay

To evaluate cognate reactivity of KSQ-001 or TIL against autologous tumor cells, KSQ-001 eTIL or TIL were co-cultured with DTCs at 1:1 ratio (100,000 cells/well of eTIL or TIL with 100,000 cells/well of DTCs) for 20-24 hours, with supernatant IFNγ level measured by MSD following manufacturer’s protocol (MesoScale Discovery, Cat# N05049A-1). HLA-ABC blocking antibody (Biolegend, Cat# 311428) and HLA-DR, DP, DQ blocking antibody (Biolegend, Cat# 361702) were included to confirm MHC dependent IFNγ release.

##### pSTAT assay

Pan-CD3 T cells, TIL, or KSQ-001 cells were thawed and rested in serum-free X-VIVO15 media (Lonza, Cat# 04-418Q) overnight. Either 100ng/mL IL-2 (PeproTech, Cat# 200-02), 100ng/mL IL-15 (CellGenix, Cat#1413-050) or 30ng/mL IL-12 (R&D system, Cat# 219IL005) was used to stimulated cells for 10-30min, 10-20min or 1hr respectively, with an excess of ice-cold PBS (Gibco, Cat#14190-136) then added to the cells. Cells were then centrifuged at 500G for 5min at 4°C and washed by ice-cold PBS.

##### In vitro A375-mOKT3 spheroid killing assay

A375-mOKT3 ‘low affinity’ cells engineered to express RFP were cultured in DMEM (Gibco, Cat #11885-084) supplemented with 10% heat-inactivated FBS (Gibco, Cat#16140-071) and 1% Pen/Strep (Gibco, Cat#15140122). 72 hours prior to assay initiation, cells were harvested via TrypLE (Gibco, Cat #12604-013), with 10,000 cells per well plated in 100μL RPMI (Gibco, Cat #11875-093) supplemented 10% heat-inactivated FBS (Gibco, Cat #16140-071) and 1% Pen/Strep (Gibco, Cat#15140122) in ultra-low attachment U-bottom plates (Corning, Cat#7007). On the day of assay initiation, TIL or KSQ-001 cells were added to spheroid plate in 100μL REP media supplemented with 6000IU/mL IL-2 (Peprotech, Cat#200-02). Images were taken via Incucyte S3 (Sartorius) at 4x magnification in the red fluorescence, brightfield, and phase channels every 6 hours for 6 days to monitor spheroid growth or regression. TIL cytotoxicity against spheroids was profiled by assessing red fluorescent intensity in each well, normalized to the first time coculture time point.

### Assessment of Editing Efficiency

#### Amplicon Sequencing (Amp-Seq)

To assess on- and off-target editing efficiencies in sgRNA/Cas9 RNP edited T cells, gDNA was extracted from edited T cells using the XTRACT16+ following the manufacturer’s protocol (Autogen, cat# KX110-96). Following the extraction of gDNA, a two-step library preparation method was performed. First step PCR consisted of a multiplex PCR reaction amplifying target sites, followed by a second step PCR adding on Illumina adapters consisting of indexing to allow for multiplexed NGS. Editing efficiency was then assessed by aligning reads to the regions of interest and the fraction of indel reads was calculated to yield a cutting score.

#### SOCS1 protein Wes

TIL or KSQ-001 cells were stimulated by ImmunoCult Human CD3/CD28/CD2 T cell activator (STEMCELL, Cat#10970) following manufacturer’s protocol overnight in REP TIL media. Cell pellets were collected, lysed by RIPA buffer (Sigma, Cat#R0278) containing Protease and Phosphatase Inhibitor (ThermoFisher, Cat# A32961). Lysates were centrifuged at 21,000g for 10min at 4°C; supernatant was subjected to BCA assay (Pierce, Cat# 23227) for protein quantification. SOCS1 (Cell Signaling, Cat# 68631S) and loading control, Vinculin (Cell Signaling, Cat# 13901S) were detected via Wes instrument (ProteinSimple) following manufacturer’s protocol.

### In vivo tumor models

For the OT1 / B16-Ova model, female C56BL/6J mice were inoculated with 0.5x10^6^ B16-Ova cells subcutaneously in the right flank. After eight days for small tumor studies (∼100mm^3^ – range of 80mm^3^-120mm^3^) or 14 days for large tumor studies (∼300mm^3^ – range of 135mm^3^-550mm^3^) mice were randomized by tumor volume and 3x10^6^ sgOlf, sgPD-1 or sgSocs1 OT-1 T cells were transferred in 0.2mL PBS via the lateral tail vein. Prior to transfer into mice, OT-1s are tested via PCR for a comprehensive list of mouse pathogens by Charles River Laboratory Testing Management. Tumor volumes were measured twice weekly using a digital caliper and tumor volume (mm3) calculated using the formula: (width2 x length)/2, where length was the longer dimension. To assess tumor growth following re-challenge in Figure 2, mice undergoing complete tumor rejection were re-challenged with 0.5x10^6^ B16-Ova cells subcutaneously on the left flank on Day 61 and monitored for tumor growth. For the PMEL / MC38-gp100 model: Female C56BL/6J mice were inoculated with 1x10^6^ MC38-gp100 cells. At the indicated tumor size, mice were randomized and received 7x10^6^ sgOlf, sgPD-1 or sgSocs1 PMEL T cells in 0.2mL PBS intravenously via the lateral tail vein. For the NOG / hIL-2 NOG engraftment model, female NOG or hIL-2 NOG mice were dosed with 10x10^6^ TIL or KSQ-001 TIL intravenously in 100μl volume. For the TIL / A375-mOKT3 model, female hIL-2 NOG mice were inoculated with 5x10^6^ A375-mOKT3 cells subcutaneously in a 50:50 mix with Matrigel (Corning Biosciences). 7 days post-inoculation, 10x10^6^ TIL or KSQ-001 were transferred intravenously.

### Tissue Processing

Single cell suspensions were generated from spleen, lymph nodes, blood and tumors. Spleens were processed using non-enzymatic digestion on GentleMacs protocol M spleen 01 01. Splenocytes were then RBC depleted with ACK (Lonza), filtered, washed, and resuspended in FACs buffer for staining. Lymph nodes were processed by gently pushing the tissue through a 35µm filter, washing and resuspending in FACs buffer for staining. Blood samples were collected by either cardiac punction at termination or via tail vein at the indicated time points and were processed using ACK (Lonza) to deplete red blood cells. Tumor tissues were dissociated using mouse/human tumor dissociation kit (Miltenyi Biotec), wherein tumors are chopped into smaller pieces, incubated for 5-10 min at 37C in the enzyme mixture and then fully dissociated using tumor program 37C_TDK_1 as recommended by manufacturer (Miltenyi Biotec).

### FACS

Single cell suspensions were processed on a 96 well plate (Falcon, Cat#353077) for flow cytometry. Cells were washed with FACS buffer (Biolegend, Cat#420201) and stained with a master mix of antibodies, viability dye (Ebioscience, Cat# 65-0865-14) and Fc block (Biolegend, Cat#422302) in FACS buffer for surface staining. Following surface staining, cells were washed by FACS buffer, fixed by fixation buffer (Biolegend, Cat#420801). For pSTAT assay, following surface staining, cells were washed by FACS buffer, fixed by Cytofix buffer (BD Biosciences, Cat#554655) and permeabilized by BD Phosflow Perm Buffer III (BD Biosciences, Cat#558058). Selected pSTAT antibody was used for pSTAT staining. A comprehensive list of antibodies used in these studies is included in the Supplementary Methods. Samples were acquired on BD Fortessa and analyzed using Flowjo software (V10, Treestar).

### CRISPR Screens

#### Human in vitro TIL expansion screen

The ‘Lib16’ sgRNA library targeting 5,137 genes with 10sgRNAs/gene and 56,408 sgRNAs total, including controls, was cloned into pKSQ017 with semi-random barcodes (see Supplementary Methods for more information on library design and lentivirus production). Lib16 includes sgRNAs targeting genes involved in T cell function, all predicted cell surface receptors, all known immune-related genes, and all genes demonstrating expression in blood.

Dissociated melanoma tumors from donor 110005746 were obtained from Conversant Bio and seeded at 1x5^6^ cells/ml in TIL-CM (RPMI 1640 + 10% HI HS, 1X HEPES, GlutaMAX, 2-Mercaptoethanol, Pen/Strep, Gentamicin) containing 300ng/ml IL-2 in a 24 well tissue culture treated plate, and incubated at 37C. Every other day, 300ng/ml IL-2 was added to cell culture assuming consumption. On Day 5, suspension cells were harvested and re-seeded at 1.5x10e6/ml in TIL-CM combined 1:1 with XVIVO-15 and supplemented with 300ng/ml IL-2, with adherent cells observed to have disappeared from the wells. Between Days 7 through Day 22, cells were expanded in IL-2 with a 1:1 ratio of TIL-CM:XVIVO-15 media supplemented with 300ng/ml IL-2 added every other day in order to maintain cell density at 1e6 cells/ml. On Day 35, cellularity was evaluated, with composition found to be 66% CD8^+^, and 32% CD4^+^. TIL5746 were frozen on Day 35 in CS10 freezing medium at 1x10^8^ cells/ml. On the first day of the screen (screen Day 0), 2x10^8^ TIL were thawed, washed in 40ml XVIVO-15 media twice, and resuspended to 2x10^6^ cells/ml in XVIVO-15 media containing 600ng/ml IL-2 and 1X DNase (Stemcell, Cat# 07900). In parallel, non-tissue culture treated 6 well plates were coated with 20ug/ml retronectin in PBS and incubated overnight in preparation for transduction. On Day 1, TIL were transduced with pKSQ017 lentivirus driving expression of Lib16 under control of the human U6 promoter, as well as driving expression of tagRFP under control of a UBC promoter. Semi-random barcodes were used to track individual clones. In parallel, an aliquot of TIL5746 were separately transduced with the KSQ041 lentivirus which drives expression of a sgRNA targeting the essential gene RPL10a under control of a human U6 promoter as well as mNeonGreen under control of a UBC promoter. Transduced cells were incubated in 6 well plates at 37C in XVIVO-15 overnight. On Day 2, TIL were combined from 6 well plates, spun at 300g for 5 minutes, supernatant removed, and TIL resuspended in complete XVIVO-15 media and rested overnight. On Day 3, transduced TIL5746 were electroporated with Cas9 mRNA (Trilink #L-7206, lot# T1COL01A) using an Amaxa 4D-Nucleofector unit and pulse code CA137 with Buffer P3 according to the manufacturer’s instructions. Following electroporation, 80μl of pre-warmed XVIVO-15 media was added to each well, mixed, and wells combined and washed. TIL were re-suspended to 1x10^6^ in complete XVIVO-15 media, and incubated overnight at 37C. On Day 4, 5x10^7^ transduced TIL5746 were frozen to determine the input sgRNA Library distribution. The TIL Rapid Expansion Phase (REP) was initiated on 25x10e6 transduced and edited TIL with a 1:200 ratio of TIL to irradiated PBMCs derived from 5 pooled donors, 600ng/ml IL-2 and 30ng/ml OKT3 using XVIVO-15 media in a 5L Grex. Cells were incubated at 37C. On Days 8, 11 and 15, 2.5L of media was aspirated from the Grex and replaced with 2.5L complete XVIVO-15 media, with 3mgs of IL-2 added to the 5L Grex. On Day 17, TIL were counted, with 13.2x10^9^ TIL5746 obtained, and analyzed for cellularity on a FACS, with 85.5% of the TIL CD8^+^, and 12.9% of the TIL CD4^+^. TIL were harvested for screen analysis by freezing three 5x10e7 cell pellets for gDNA preparations. The sgRNA distribution was compared between TIL harvested on Day 4 and Day 17.

#### In vivo Cas9-Tg x TCR-Tg CD8 T cell syngeneic tumor screens

The ‘Lib30’ sgRNA library targets 369 genes involved in T cell function with 10sgRNAs per gene and 3089 sgRNAs total, including controls, was cloned into pKSQ044. Lib31 sgRNA library targets 1004 predicted CSR genes with 8 sgRNAs/gene and 11,148 sgRNAs total, including controls, was also cloned into pKSQ044 (see Supplementary Methods for more information on library design and lentivirus production). Cas9-Tg x TCR-Tg OT1 or PMEL CD8 T cells were isolated from freshly harvested mouse spleens and dissociated using a GentleMACS (Miltenyi), and with CD8 T cells purified by negative selection (EasySep Mouse CD8+ T cell isolation kit). CD8s were then placed in T225 flasks at a concentration of 1M CD8s/ml and activated with CD3/CD28 Dynabeads and 2ng/ml mouse rIL-2. The following day, T cells were transduced wither either lentivirus (OT1 B16-Ova screen) or retrovirus (PMEL / MC38-gp100 screen). Dynabeads were removed the following day, with cells resuspended on cRPMI + 32ng/ml IL-2. On Day 4, transduced T cells were harvested and selected by positive selection using either Thy1.1 for OT1s or hCD2 for PMELs. 5x10^6^ Thy1.1^+^ Cas9-Tg x OT1 CD8 T cells or 7x10^6^ hCD2^+^ Cas9-Tg x PMEL CD8 T cells were injected into tumor-bearing mice i.v. via tail vein in 200μL PBS, with 10x10^6^ CD8^+^ T cells saved to determine the sgRNA distribution of the input population of T cells. Group sizes were 7-8 mice per library. At the indicated time following adoptive transfer of cells, mice were euthanized and blood was harvested in EDTA tubes, and stored at -80C. Tumors, spleens, tumor draining and non-draining lymph nodes were harvested and processed further for extraction of CD8 T cells using CD8a Microbeads (Miltenyi, cat# 130-049-401) from the spleen and CD45 Microbeads (Miltenyi, cat# 130-052-301) from the tumor. Tumors were digested using the Miltenyi Tumor Dissociation Kit (Cat# 130-096-730) according to the manufacturer’s instructions. Genomic DNA was isolated using the Qiamp Blood Midi and Maxi kids, according to the manufacturer’s instructions.

#### SOCS1 sgRNA Tiling Screen

Primary human CD8^+^ T cells were isolated from PBMCs, with ∼300x10^6^ CD8 T cells obtained by negative selection using a EasySep Human CD8+ T cell Isolation Kit (StemCell; Cat# 17953). T cells were activated with Immunocult Human CD3/CD28/CD2 T cell activators (Stemcell; Cat# 10990) in the presence of 10U human IL-2 in X-VIVO-15 media. On the following day, CD8 T cells were harvested, counted, and transduced with lentivirus expressing 134 sgRNAs targeting genomic positions across the full length of the SOCS1 gene as well as multi-cutter sgRNAs as depleting and Olfactory genes as neutral controls. Two days after transduction, T cells were electroporated with Cas9 mRNA (Trilink) and expanded inn 10U human IL-2 in X-VIVO-15 media for 10 additional days. Cells were then harvested; DNA was extracted and amplicons spanning the lentiviral genomic regions containing the sgRNA cassettes in the library were amplified by polymerase chain reaction (PCR) and sequenced by next-generation sequencing (NGS).

#### CRISPR Screen Pre-processing

Counts of sgRNA frequency were generated from FASTQ sequencing files by counting the number of occurrences of each 20nt sequence READ1. For libraries that employed a unique molecular identifier the frequency of each UMI was generated by tabulating the number of occurrences of each 12nt prefix of READ2. For libraries containing UMIs the resulting count tables were further cleaned by requiring 1) the UMI matches the mixing code used to synthesize the library 2) does not contain any ambiguous/uncalled bases 3) is not potential index-bleed (exactly matching a more abundant guide/UMI on the same sequencing run). To minimize the impact of singleton/low frequency clones a two-component mixture model was fit to the count frequency distribution and guides present in the low abundance component were dropped.

#### Analysis of in vitro TIL CRISPR Screens

To identify hits from the screen we calculated the fold-enrichment of each candidate gene in an “end-point” relative to a “reference” sample. The reads from raw FASTQ files were counted, and the number reads representing each guide were tabulated. For each guide, a log-ratio between the screen end-point and the reference sample was calculated. To compute a robust gene-level enrichment score, guide level scores were aggregated by taking the median enrichment score across all guides. We assigned each guide a conservative p-value equal to the percentile of its logFC among all guides in the library. To calculate a p-value for gene enrichment individual guide-level p-values were combined using Fisher’s method. The resulting p-values were adjusted for false discovery rates by the Benjamini-Hochberg method. To standardize (Z-score) the resulting scores, logFC scores were centered and scaled by subtracting the median enrichment score and dividing by the median absolute deviation to calculate an overall effect size for each gene.

#### Analysis of in vivo B16-OVA and MC38-gp100 CRISPR Screens

MAGeCK-mle(Version 0.5.9.3) was used to perform CRISPR screen analysis. Genes with guide numbers fewer than 5 were removed from the analysis. The number of unique UMI for each sgRNA were summarized as input clone counts. Clone counts were organized from all in-vivo tumor samples and input samples into a count matrix. Due to the large number of zero counts in the matrix, an altered RLE normalization was performed on raw count matrix by using geometric mean of gene and size factor of one sample calculated by only non-zero counts. Normalized count matrix was used as MAGeCK-mle input. Design matrix was composed of two columns: input column with all “1” as baseline level, and tumor column with only tumor samples as “1” and the rest samples as “0”. Gene level beta values from MAGeCK-mle output were used to evaluate the effect size of knocking out the corresponding gene.

### scRNA-Seq

Droplet-base 5’ single-cell RNA sequencing (scRNA-Seq) was performed by the 10x Genomics platform and libraries were prepared by the Chromium Single Cell 5’ Reagent kit according to the manufacturer’s protocol (10x Genomics, CA, USA). The Cell Ranger (version 6.0.1) was used for gene expression quantification, TCR sequence assembly, and cell identification. Cell level quality control was performed using the function quickPerCellQC() from scater (version 1.18.6). Per-cell size-factor normalization was performed using the computeSumFactors() function from scran (version 1.18.7). Cell lineages were annotated automatically using clustifyr (version 1.2.0) and the Haemosphere mouse RNA-seq database(58). T cells were isolated by computational gating based on expression of Cd3d, Cd3e, and Cd3g. Seurat (version 4.1.0) was used to identify clusters and perform differential gene expression analysis. Differentially expressed genes were identified using the Wilcoxon Rank Sum test. The hypergeometric test was performed to assess treatment group enrichment in each of the identified clusters. ProjecTILs (version 2.0) was used to map our data onto scRNA-Seq data from Miller et al (45). *F*gsea (version 1.26.0) was used to perform GSEA on the CD8 expressing T cells using Tex gene signatures from Miller *et al.* and Beltra(45, 59). GSVA (version 1.48.2) was used to perform GSVA on the CD8 expressing T cells using Tex gene signatures from Miller *et al.* and Beltra *et al.* The extent of clonal expansion per cluster was quantified using the StartracDiversity() function from scRepertoire (version 1.10.0).OT1 T cells were identified by searching the TCR repertoire for cells containing both the OT1 CDR3 Tcra amino acid sequence (CAASDNYQLIW) and the OT1 CD3R Tcrb amino acid sequence (CASSRANYEQYF).

### Statistical Analyses

Data from mouse efficacy studies were expressed as mean +/- SEM. Statistical significance between groups was determined by either 2-way ANOVA or a two-tailed unpaired Student’s *t* test as indicated. Statistical analyses were performed using GraphPad prism software, with ns = no significance; * = p < 0.05; ** = p < 0.01; *** = p < 0.001 and **** = p < 0.0001. Simpson Diversity Index was used to evaluate the diversity of TCRs within a population of TIL by measuring the probability that two TCRs taken randomly from a population of TCRs will be different TCR sequences, with the higher the Simpson Diversity Index value, the greater TCR diversity in a population of TIL. The Morisita Index was used to evaluate TCR overlap between two populations of TIL by quantifying how similar two TCR repertoire samples as determined by TCR-Seq are in terms of the overlapping TCR sequences and how well represented those sequences are within the two populations. Statistical significance between treatment groups in scRNA-Seq clusters depicted in Figure 3h was determined by hypergeometric test.

### Study Approval

All procedures involving the care and use of animals were reviewed and approved by the Institutional Animal Care and Use Committee (IACUC) and were conducted at the AAALAC accredited CRADL under protocol 2021-1252 in accordance with associated regulations and guidelines.

### Data Availability

The T cell CRISPR screening data from Figure 1 and scRNA-Seq data from Figure 3 are available in Supplementary Tables 1-3, with data also uploaded and available at GEO (GSE237695). Values for data points described in Figures 2, 3, 6, 7 as well as Figures S1, S2, S3, S6 and S7 can be found in the supporting data values file.

## Supporting information

Supporting Data Values

Table S3 PMEL MC38gp100 Screen

Table S2 OT1 B16Ova Screen

Table S1 TIL Screen

## Author Contributions

M.R.S, K.W., F.A.S, F.S., L.C. and M.J.B. conceived of experiments. C.W., S.S., S.L., C.C., Z.S., T.X., H.G., L.W., F.T., P.D., R.A.L, T.E.G., N.C., A.F.H, N.T, C.B, I.L.M, K.S., J.J.M, S.K., and G.V.K. conducted experiments and analyzed data. M.J.B. wrote the manuscript. M.R.S., S.S., C.D., F.S., L.C. and M.J.B provided supervision. Co-first authorship determined as M.S., C.W., Z.C., S.S. and S.L. served as scientific leads for functional genomics, in vivo pharmacology, computational biology and in vitro pharmacology functions, respectively.

## Acknowledgements

We would like to thank Dr. Shari Pilon-Thomas, Dr Michal Besser and Dr Gustavo Martinez for input and thoughtful comments on the manuscript.

## Notes

Conflict-of-interest statement: All authors are current or former employees and shareholders of KSQ Therapeutics

### Competing Interest Statement

All authors are current or former employees and shareholders of KSQ Therapeutics

