## Supplementary Information for "Rational design of a SOCS1-edited tumor infiltrating lymphocyte therapy for solid tumors using CRISPR/Cas9 screens"

**For**

**Contents**


- **Supplementary Methods**
  - Design and generation of screening libraries
  - sgRNA library virus production
  - T cell transduction
  - GUIDE-seq
  - Engineering of high- and low-affinity A375-mOKT3 cells
  - sgRNA sequences
  - Amplicon sequencing primers for on- and off-target editing assessment
  - Antibody Table

- **Supplemental Data**
  - Supplemental Data 1: Human TIL screen QC
  - Supplemental Data 2: OT1 / B16-Ova screen QC
  - Supplemental Data 3: PMEL / MC38-gp100 screen QC

- **Supplementary Figures**
  - **Figure S1:** Development of a syngeneic tumor models sensitive and insensitive to inactivation of PD-1 in adoptively transferred TCR-Tg CD8 T cells
  - **Figure S2:** Inactivation of SOCS1 enhances the in vivo anti-tumor potency of CD8 T cells and drives the accumulation of $CD44^{+}CD62L^{+}$ $T_{cm}$ memory cells in blood
  - **Figure S3:** FACS analysis of innate cell subsets from sgOlf, sgSocs1 and sgPD-1 OT1s from the TME 7 days following transfer
  - **Figure S4:** scRNA-Seq of sgOlf, sgSocs1 and sgPD-1 OT1s from the TME 7 days following transfer
  - **Figure S5:** Identification of the genomic locations of u728 sgRNA off-target cut sites
  - **Figure S6:** Characterization of KSQ-001
  - **Figure S7:** Sensitivity of KSQ-001, TIL, and sgSOCS1-edited $CD3^{+}$ T cells to IL-2 and IL-15

### Supplemental Methods

#### Design and generation of screening libraries:

Viral vectors were constructed for both human and mouse T-cell screening. Human libraries were in a pLenti6-based lentiviral backbone that contained a human U6 promoter cassette for guide cloning, as well as a puromycin resistance marker and tagRFP fluorescent marker (pKSQ017) or a Thy1.1 selectable marker (pKSQ144). For retrovirus, the vector was a self-inactivating murine stem cell virus (MSCV) backbone containing the same human U6 promoter guide cloning cassette as well as a human CD2 marker for cell surface expression and bead enrichment of infected cells. The vectors used for TCR-Tg CRISPR screens additionally contained a library of randomly synthesized 12-mer barcodes with the nucleotide sequence BVHDBVHDBVHD (IUPAC mixed base codes). These random barcodes (theoretical diversity of 531,441 unique sequences), when randomly paired with the guide sequence during cloning, allow the ability to independently trace millions of unique infection events in immune cells.

Libraries were generated by searching for PAM sites (NGG) in the human and mouse genome that were expected to correspond to sgRNAs that would cut the coding sequence of genes of interest. sgRNAs were chosen for the library that were predicted to cut the genome in at most one site, while not overlapping with known high prevalence SNPs and having few (<10) closely related sites in the genome that could represent likely off-target sites. Several classes of controls were included, non-cutting controls, olfactory receptor cutting controls not expected to impact biology in immune cells, lethal shredder controls that cut the genome thousands of times, and (for human only) fingerprinting controls that cut high-prevalence SNPs in human essential genes and allowed internal identification of samples by donor. The final set of filtered sgRNAs were then synthesized on a microarray (Agilent oligo library synthesis) as oligonucleotides with adapters, PCR amplified, and cloned using the type IIs restriction enzyme BbsI into the final lentiviral or retroviral vector.

#### sgRNA library virus production

Lentivirus production: Lentivirus was generated by lipid transfection of packaging plasmids into HEK293T cells. For a 10-layer CellStack (Corning), 300 million HEK239T cells were plated in 1 L of DMEM + 10% FBS. 24 Hours later transfection was performed using 461 ug of sgRNA library plasmid, 231 ug of packaging plasmid psPax2 (Gag-Pol), and 115 ug of pMD2.G packaging plasmid (VSV-G). Plasmid DNA was added to 37.77 ml of room temperature OptiMEM media. 2428 ul of TransIT transfection reagent (Mirus Bio) was added, mixed by vortexing, and incubated for 20 minutes before proceeding. OptiMEM+DNA+TransIT mixture was added to 1 liter of DMEM + 10% FBS. Media was aspirated from HEK293T cells and fresh media containing transfection mixture was added. 24 hours after transfection the media was replaced 1 liter of UltraCulture media (Lonza) supplemented with L-glutamine. 48 hours after transfection the supernatant was harvested from the cells, incubated for 1 hour at 37C in the presence of 50 units/ml benzonase, filtered using a 0.45 uM bottle top PES filter, and concentrated by tangential flow filtration on a spectrum labs KrosFlow mPES hollow fiber filter (100kD molecular weight cutoff). Virus was concentrated approximately 10-50x (depending on initial volumes being processed).

Retroviral Production: Retrovirus was generated by transient transfection of Phoenix-Eco retroviral packaging cells with library plasmid pools. 5 million Phoenix-Eco cells were plated per 10 dish 24 hours before transfection in DMEM + 10% FBS. Transfection is carried out by mixing 10ug of library plasmid pool with 327 ul optiMEM media. 21 ul of Mirus TransIT-293 transfection reagent is added to this mixture, vortexed briefly, and incubated for 20 mins. Transfection mixture is then added dropwise to previously plated phoenix cells and incubated. For infection of mouse CD8 cells, media is replaced with 5ml of RPMIc (RPMI + 10% HI FBS, 20 mM HEPES, 50 uM 2-mercaptoethanol). Virus is frozen and titered for subsequent large-scale transduction.

-

**T cell Transduction**

TIL: To transduce TIL5746, retronectin-coated non-TC-treated 6 well plates were washed with 1ml PBS, with 200ml of pKSQ017 Lib16 or RPL10a virus added and spun at 2000g for 2hrs. $5x10^6$ TIL/well in 1ml complete XVIVO-15 media were added to each well (total volume 1.2ml) so that 200 x10e6 TIL5746 were transduced with Lib16, and 10x10e6 TIL5746 were transduced with sgRNAs targeting RPL10a. Plates were spun at 1000g for 10 minutes, and then transferred to 37C for 3 hours, followed by addition of 3ml of complete XVIVO-15 to all wells, and incubation at 37C overnight.

Mouse TCR-Tg cells: To transduce mouse CD8 TCR-Tg T cells isolated from spleens and activated with Dynabeads and 4ng/ml IL-2 on Day 0, non-TC-treated 6 well plates were coated in parallel with retronectin overnight at 4C. On Day 1, plates were washed with 1ml PBS and blocked with DPBS + 2% BSA for 30 minutes. Activated mouse TCR-Tg CD8 cells were then harvested and resuspended at a concentration of $1.5x10^6$/ml in thawed concentrated lentivirus or retrovirus supernatant with 2ng/ml IL-2 and 5mg/ml protamine sulfate. 2ml of virus + CD8 mixture is pipetted in each well of the retronectin-coated 6-well plate, with plates centrifuged at 600g for 90 mins, and then transferred to a 37C TC incubator for 5 hours. 2ml of cRPMI + 2ng/mL IL-2 was then added to each well and incubated overnight at 37C. Cells were harvested, washed, and re-suspended in cRPMI + IL2 for further expansion on the following day. The lentivirus (Thy1.1) and retroviruses (CD2) used for the OT1 and PMEL transductions, respectively, contain selectable markers for positive selection, with this selection step occurring on Day 3 following transduction. On Day 4, cells were cryopreserved for future use.

**GUIDE-seq**

The Guide-seq protocol was modified from Tsai et al. 2015(1) and Nagendra Palani (https://www.protocols.io/view/guide-seq-simplified-library-preparation-protocol-kxygxmwool8j/v1 ). The double-strand oligodeoxynucleotide (dsODN) was formed by annealing the following modified single-strand DNA:

5′-Phos-G*T*TTAATTGAGTTGTCATATGTTAATAACGGT*A*T-3′ and
5′-Phos-A*T*ACCGTTATTAACATATGACAACTCAATTAA*A*C-3′.

Each oligonucleotide is phosphorylated at the 5’ end and includes phosphorothioate bonds (noted by asterisks). For GUIDE-Seq, primary T cells were electroporated with RNP complexes as described above with the addition of 200pmol of the annealed dsODN. The single-tail Y adapter was formed after annealing the following DNA strands: CTACAAGAGCGGTGAGT and 5’-Phos-CTCACCGCTCTTGTAGSNNNNNNNNCTGTCTCTTATACACATCTCCGAG*C. The tail of the adapter includes a unique molecular identifier (indicated by the sequence SNNNNNNNN). Double-stranded genomic DNA was quantified by the Qubit Broad Range Assay (Cat # Q32853) and 1 µg of DNA from each sample was used for library preparation using the NEBNext Ultra II FS DNA Library Prep kit (Cat # E6177L). Genomic DNA was fragmented to 300-700bp (37°C incubation for 10 minutes) and 5 µl of 10 µM annealed adapter was used in the subsequent adapter ligation step. Purified adapter ligated DNA was used for two separate touch-down PCR enrichment reactions of dsODN containing fragments (plus and minus strand orientations each paired with a primer binding to the Y adapter). These primers are:

Minus strand dsODN primer –
TCGTCGGCAGCGTCAGATGTGTATAAGAGACAGNNNATACCGTTATTAACATATGACAACTCAATTAAA
*C,
Plus strand dsODN primer – TCGTCGGCAGCGTCAGATGTGTATAAGAGACAGNNNGTTTAATT
GAGTTGTCATATGTTAATAACGGTA*T,
and Adapter primer – GTCTCGTGGGCTCGGAGATGTGTATA AGAGACA*G.

The double-strand DNA products from these reactions were quantified and 5ng from each sample’s plus and minus reactions were pooled for indexing PCR using primers from the Nextera DNA CD Indexes 96 well plate (Cat # 20018708). Final PCR products were quantified and library distribution was measured by D1000 TapeStation (Cat # 5067-5582 and 5067-5583). All samples were pooled by equal mass and purified by 1.2X SPRI bead clean-up. The concentration of the final pool was quantified by Kapa Illumina Library quantification kit (Cat #KK4923). Final pools were loaded onto the NextSeq with at least 10% PhiX spike-in to ensure sequence diversity.

**Engineering of high- and low- affinity A375-mOKT3 cells**

The A375-mOKT3 cell line was generated by infecting A375 melanoma cells with a lentiviral vector encoding a blasticidin resistance gene and either a high affinity (parental OKT3 sequence) or low affinity version (two amino acid substitutions to reduce affinity for CD3 ~1000 fold) of the mOKT3 protein.(2) Individual clones were selected and tested for their ability to stimulate TIL in vitro.

High affinity OKT3:
MERHWIFLLLLSVTAGVHSQVQLQQSGAELARPGASVKMSCKASGYTFTRYTMHWVKQRPGQGLEWIGYI
NPSRGYTNYNQKFKDKATLTTDKSSSTAYMQLSSLTSEDSAVYYCARYYDDHYCLDYWGQGTTLTVSSGGG
GSGGGGSGGGGSQIVLTQSPAIMSASPGEKVTMTCSASSSVSYMNWYQQKSGTSPKRWIYDTSKLASGVP
AHFRGSGSGTSYSLTISGMEAEDAATYYCQQWSSNPFTFGSGTKLEINSHFVPVFLPAKPTTTPAPRPPTPAP

TIASQPLSLRPEACRPAAGGAVHTRGLDFACDIYIWAPLAGTCGVLLLSLVITLYCNHRNRRRVCKCPRPVVKS
GDKPSLSARYV

Low affinity OKT3[LT]:
MERHWIFLLLLSVTAGVHSQVQLQQSGAELARPGASVKMSCKASGYTFTRYTMHWVKQRPGQGLEWIGYI
NPSLGTTNYNQKFKDKATLTTDKSSSTAYMQLSSLTSEDSAVYYCARYYDDHYCLDYWGQGTTLTVSSGGGG
SGGGGSGGGGSQIVLTQSPAIMSASPGEKVTMTCSASSSVSYMNWYQQKSGTSPKRWIYDTSKLASGVPA
HFRGSGSGTSYSLTISGMEAEDAATYYCQQWSSNPFTFGSGTKLEINSHFVPVFLPAKPTTTPAPRPPTPAPTI
ASQPLSLRPEACRPAAGGAVHTRGLDFACDIYIWAPLAGTCGVLLLSLVITLYCNHRNRRRVCKCPRPVVKSG
DKPSLSARYV

**sgRNA sequences**

| sgRNA | Description : Species : Target | Sequence |
|---|---|---|
| k1vc | sgOlf : Mouse : Olfactory receptor | GGAAGAAGTACATCTGCAAG |
| 336b | sgPD-1 : Mouse : PD-1 | CGGAGGATCTTATGCTGAAC |
| 4u9j | sgSocs1: Mouse SOCS1 | GCCGGCCGCTTCCACTTGGA |
| u728 | Therapeutic sgRNA : Human : SOCS1 | GACGCCTGCGGATTCTACTG |
| a7mm | Control sgRNA: Human ORA1 | GCTGACCAGTAACTCCCAGG |
| anvh | Candidate sgRNA : Human : SOCS1 | GGCCGGCCTGAAAGTGCACG |
| v2v0 | Candidate sgRNA : Human : SOCS1 | GCGGCTGCGCGCCGAGCCCG |
| frmu | Candidate sgRNA : Human : SOCS1 | CGCACCAGGAAGGTGCCCAC |
| xsri | Candidate sgRNA : Human : SOCS1 | GGACGCCTGCGGATTCTACT |
| kntc | Candidate sgRNA : Human : SOCS1 | GGCTGCCATCCAGGTGAAAG |
| 8usj | Candidate sgRNA : Human : SOCS1 | GCCGGCCGCTTTCACCTGGA |
| erv7 | Candidate sgRNA : Human : SOCS1 | CTTAGCGTGAAGATGGCCTC |
| 7bv8 | Candidate sgRNA : Human : SOCS1 | AGCGCGCTCCTGGACGCCTG |
| qd5u | Candidate sgRNA : Human : SOCS1 | TGGACGCCTGCGGATTCTAC |
| kipc | Candidate sgRNA : Human : SOCS1 | AGTGCTCCAGCAGCTCGAAG |

| | | |
|---|---|---|
| i1d4 | Candidate sgRNA : Human : SOCS1 | ACGCCTGCGGATTCTACTGG |
| CCR5 R-30 | Control sgRNA : Human : CCR5 | GTGTTCATCTTTGGTTTTGT |
| CCR5 R-25 | Control sgRNA : Human : CCR5 | GTAGAGCGGAGGCAGGAGGC |

**Amplicon sequencing primers for on- and off-target editing assessment**

| sgRNA | 3'-Primer | 5'-Primer |
|---|---|---|
| k1vc | TCAATAGCCATGACAGTCAGAAG | ACAGCTACCATACCTAAGATGCT |
| 336b | TATGATCTGGAAGCGGGCAT | CAAATGCCACCTTCACCTGC |
| 4u9j | CTTCTTGGTGCGCGACAG | GTCACGGAGTACCGGGTTAAG |
| u728 | CACGCACTTCCGCACATT | GAGGCCATCTTCACGCTAAG |
| a7mm | CAATGTGATGGGGTAAATGAACA | GATATTCCTCTCCCCTTTCATG |
| anvh | ATGCGAGCCAGGTTCTCG | CTTAGCGTGAAGATGGCCTC |
| v2v0 | GAGGCCATCTTCACGCTAAG | CACGCACTTCCGCACATT |
| frmu | GAAGAGGCAGTCGAAGCTCT | CACGCACTTCCGCACATT |
| xsri | CACGCACTTCCGCACATT | GAGGCCATCTTCACGCTAAG |
| kntc | GTGAAGATGGCCTCGGGAC | ATGCGAGCCAGGTTCTCG |
| 8usj | ATGCGAGCCAGGTTCTCG | CTTAGCGTGAAGATGGCCTC |
| erv7 | GAAGAGGCAGTCGAAGCTCT | CTTAGCGTGAAGATGGCCTC |
| 7bv8 | GAGGCCATCTTCACGCTAAG | CACGCACTTCCGCACATT |
| qd5u | GAGGCCATCTTCACGCTAAG | CACGCACTTCCGCACATT |
| kipc | CTTAGCGTGAAGATGGCCTC | ATGCGAGCCAGGTTCTCG |
| i1d4 | CACGCACTTCCGCACATT | GAGGCCATCTTCACGCTAAG |

| | | |
|---|---|---|
| CCR5 R-30 | CAGGGTGGAACAAGATGGATTA | GTTGAGCAGGTAGATGTCAGTC |
| CCR5 R-25 | CAGGGTGGAACAAGATGGATTA | GTTGAGCAGGTAGATGTCAGTC |
| U728_ON | GTCAGATGTGTATAAGAGACAGGGCGCGCAGCCGCTCGTGCG | CGGAGATGTGTATAAGAGACAGATTCCGTTCGCACGCCGATTACCGG |
| OT1 | GTCAGATGTGTATAAGAGACAGATTATTTACTGTCTTGCTCCAGGGCTA | CGGAGATGTGTATAAGAGACAGCATTCTCACCAGAGCCCATTAGTATGA |
| OT2 | GTCAGATGTGTATAAGAGACAGACCCCATCTGCTCTGGCCCACA | CGGAGATGTGTATAAGAGACAGTGCCATAGAACAGACTCTAACACCAGCT |
| OT3 | GTCAGATGTGTATAAGAGACAGAGTTTCCAGCCAGGACAGCCCCACA | CGGAGATGTGTATAAGAGACAGATCTCACTCACCCAGTGGGAAGGG |
| OT4 | GTCAGATGTGTATAAGAGACAGCACACACACACACGCACACCCTATCATA | CGGAGATGTGTATAAGAGACAGATTGTAGCCTCATATTTGCCCCCACG |
| OT5 | GTCAGATGTGTATAAGAGACAGCTGCTCTATTTGTTACTCACCTGGGA | CGGAGATGTGTATAAGAGACAGACACTGCCATATTCAAGCAGATTAAGTA |
| OT6 | GTCAGATGTGTATAAGAGACAGATTGCAATCTCAAATTTCTGGCCTCAA | CGGAGATGTGTATAAGAGACAGGTGAGATCCCGTCTGTGCAAAAAATTTA |
| OT7 | GTCAGATGTGTATAAGAGACAGGGAAGTCCGGCTCCCTCTCCACATTC | CGGAGATGTGTATAAGAGACAGGCTCATCTGGCAGCCAGGACA |
| OT8 | GTCAGATGTGTATAAGAGACAGCCAGCATGCTTCTTTCCACCACA | CGGAGATGTGTATAAGAGACAGTGATGATGCAGCTCCTCCTCAAAGGG |
| OT9 | GTCAGATGTGTATAAGAGACAGCATGACTAACAAGCTGTGAAAGTACTCA | CGGAGATGTGTATAAGAGACAGAAGTGCTTGGTTTAATGGGAAAAGAAAA |
| OT10 | GTCAGATGTGTATAAGAGACAGCAACACCTTGAACTAACATTACTAGCTA | CGGAGATGTGTATAAGAGACAGGGCTGTGATATTAAAGAGTAGTCCCTAA |
| OT11 | GTCAGATGTGTATAAGAGACAGGGCAGTCATGATCAAGTCAATCCCTTG | CGGAGATGTGTATAAGAGACAGATGGAACCTTGCTATGTTGCCTAGGG |
| OT12 | GTCAGATGTGTATAAGAGACAGCAGCAGCTGCTTTCTGTCCTCTCTTTA | CGGAGATGTGTATAAGAGACAGAGGCTCAATGATTCAAGGCCTGCT |
| OT13 | GTCAGATGTGTATAAGAGACAGACCACCTTGATCTTGTTGCTCTGGG | CGGAGATGTGTATAAGAGACAGTCAAGAGTCTGGAGTGAGGGCCTCATG |
| OT14 | GTCAGATGTGTATAAGAGACAGGTAGCCAAAAGTCAGTTACTACATCTCT | CGGAGATGTGTATAAGAGACAGAGACCCAAGCTTGGAAAGTGATCTTTG |
| OT15 | GTCAGATGTGTATAAGAGACAGTGTCTGGGTGCTCTGCCCAATGCCCTGA | CGGAGATGTGTATAAGAGACAGCCTGCTCTCCCTAACAGCAGCA |
| OT16 | GTCAGATGTGTATAAGAGACAGTGTGGGACCAATGGCTGGGCAA | CGGAGATGTGTATAAGAGACAGAAGATCTCAGCCCACCCCTAGCACCA |
| OT17 | GTCAGATGTGTATAAGAGACAGAGTTCTAGCTTAGAAGCTCATTGCTCT | CGGAGATGTGTATAAGAGACAGCCTGTTTATCCTGAGAGCCTCTGCTTC |
| OT18 | GTCAGATGTGTATAAGAGACAGCCAGATACCCTTACCCTGAATACTCTGC | CGGAGATGTGTATAAGAGACAGCCCATGCTGCCCATATGGCTGTAATTA |

| | | |
|---|---|---|
| OT19 | GTCAGATGTGTATAAGAGACAGGCACTA CCAACCTGAAGGTGTGGCT | CGGAGATGTGTATAAGAGACAGCCTCCC GGTGAGCCTGGTGTT |
| OT20 | GTCAGATGTGTATAAGAGACAGCATCCT CTTCTGAGAACCAACAATTCCA | CGGAGATGTGTATAAGAGACAGCCATCG GGTCCAGCCACCA |
| OT21 | GTCAGATGTGTATAAGAGACAGATCAGC AAGCAAGAGCCTTCCTGG | CGGAGATGTGTATAAGAGACAGGGCTGA GAGGCATTGCTGACG |
| OT22 | GTCAGATGTGTATAAGAGACAGCGGCCT GCCTGCGACCCGAGA | CGGAGATGTGTATAAGAGACAGCTCCAC TGCACCTCGCCAGA |

### FACS Antibodies

| Antigen | Color | Vendor | Cat # | Antigen species | Figures |
|---|---|---|---|---|---|
| CD4 | PerCP Cy5.5 | BD Bioscience | 561115 | Mouse | 2 |
| CD8 | APC | BD Bioscience | 553035 | Mouse | 2 |
| Va2 | PE | BD Bioscience | 553289 | Mouse | 2 |
| CD45 | BUV661 | BD Bioscience | 612975 | Mouse | 3 |
| CD3 | BV650 | BD Bioscience | 564378 | Mouse | 3 |
| CD8 | BUV395 | BD Bioscience | 563786 | Mouse | 3 |
| CD4 | BUV496 | BD Bioscience | 612952 | Mouse | 3 |
| Vb5.1/2 | BV421 | BD Bioscience | 742998 | Mouse | 2, 3 |
| Va2 | BB700 | BD Bioscience | 746041 | Mouse | 3 |
| Slamf-6 | APC | Biolegend | 134610 | Mouse | 3 |
| Foxp3 | PE | eBio | 12-5773-82 | Mouse | 3 |
| CD44 | BUV737 | BD Bioscience | 612799 | Mouse | 2, 3 |
| CD39 | PE.Cy7 | Biolegend | 143806 | Mouse | 3 |
| CD62L | BV605 | BD Bioscience | 563252 | Mouse | 2, 3 |
| PD-1 | FITC | Biolegend | 135214 | Mouse | 3 |
| CD45 | BV786 | BD Bioscience | 564225 | Mouse | S3 |
| CD11b | BUV737 | BD Bioscience | 612800 | Mouse | S3 |
| CD11c | PE | Biolegend | 117308 | Mouse | S3 |
| Ly6C | FITC | Biolegend | 128006 | Mouse | S3 |
| Ly6G | BB700 | BD Bioscience | 566435 | Mouse | S3 |
| NKp46 | APC | Biolegend | 137608 | Mouse | S3 |
| CD103 | BV421 | BD Bioscience | 562771 | Mouse | S3 |
| MHC-II | BV605 | BD Bioscience | 563413 | Mouse | S3 |
| F4/80 | BUV395 | BD Bioscience | 565614 | Mouse | S3 |
| CD115 | BV650 | BD Bioscience | 750890 | Mouse | S3 |
| pSTAT5 (pY694) | PE | BD Bioscience | 612567 | Human | 6 |
| pSTAT4 (pY693) | PE | BD Bioscience | 558249 | Human | 6 |
| CCR7 | Alx 647 | BD Bioscience | 560816 | Human | S6 |
| CD45RA | PerCP Cy5.5 | BD Bioscience | 563429 | Human | S6 |
| CD45RO | PE Cy7 | BD Bioscience | 560608 | Human | S6 |
| Live/Dead | e780 | ebioscience | 65-0865-14 | N/A | All FACS experiments |
| CD45 | BV510 | BD Bioscience | 563204 | Human | S6 |
| MCSP | Alx 647 | BD Bioscience | 562414 | 9.2.27 | S6 |
| CD3 | BV421 | BD Bioscience | 563798 | SK7 | S6 |
| CD25 | BUV661 | BD Bioscience | 741685 | Human | S6 |

| | | | | | |
|---|---|---|---|---|---|
| CD4 | Alx 700 | BD Bioscience | 560836 | Human | S6E |
| CD8 | BUV395 | BD Bioscience | 563795 | Human | S6 |
| IFNg | PE-Cy7 | BD Bioscience | 560741 | Human | S6 |
| TNFa | Alx488 | BD Bioscience | 557722 | Human | S6 |
| IL-2 | PE | BD Bioscience | 559334 | Human | S6 |

**Supplemental Data**

Supplemental Data 1: Human TIL Screen QC

Background: This screen investigated the effect of CRISPR knockout on human TIL using lib16, a sub-genome library. The following table describes the two samples that were compared.

1. Results: The screen analysis was performed on zorya. A QC report was generated with clean_count output saved.

   1.1: Description of samples included in analysis:

| Library | Day of Analysis | Number of Samples |
|---|---|---|
| Lib16 | Day 4 | 1 |
| Lib16 | Day 17 | 1 |

2. Library/Screen QC

2.1: Screen sequencing purity: What fraction of sequenced reads match to the expected library design?

Recovered sgRNAs: 85%. Missing: 15%.

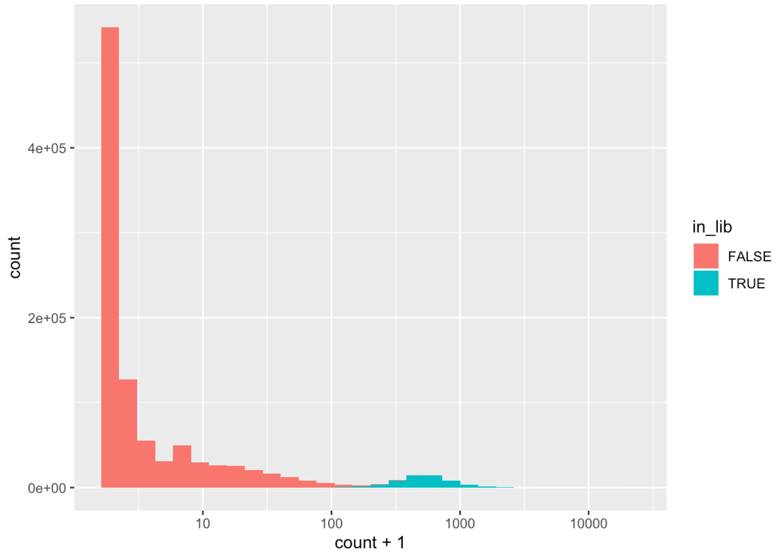


The libraries look as expected, majority of reads match to the library design, and the sequences matching the library design have much higher counts than sequences that are not part of the library design.

2.2: Screen Coverage

We expect the majority of the sgRNA library to be recovered.

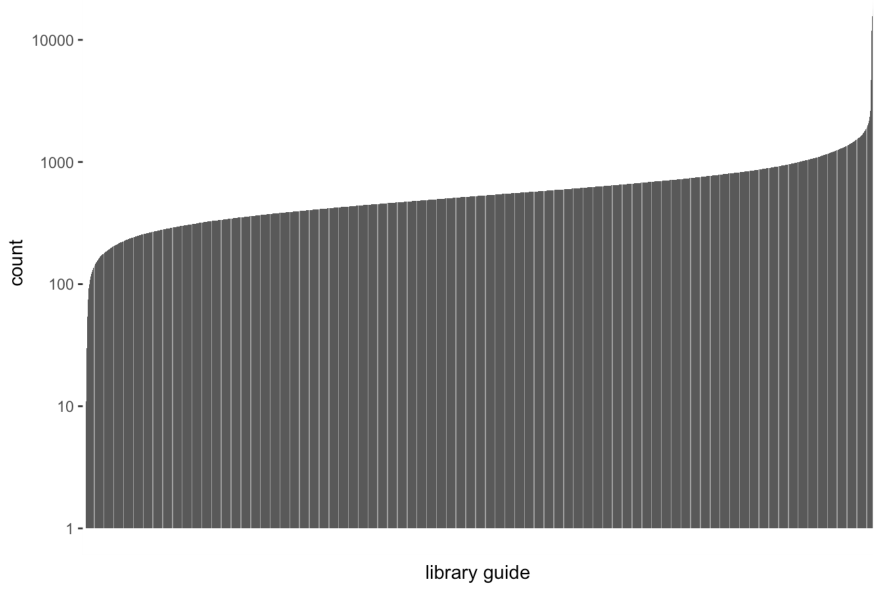


Majority of guides have uniform counts.

2.3: Gene Level QC

2.3.1: Negative controls should drop out

lib16 included controls targeting multiple positions across the genome (type="control_multicutter"). We expect these guides to drop out compared to the average guide in the library.

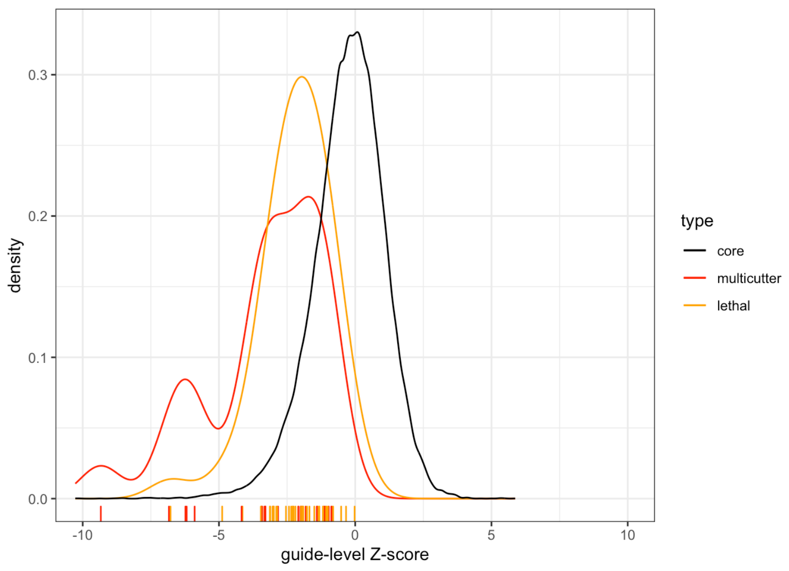

Control guides targeting essential genes or high-copy regions of the genome drop out, confirming Cas9 activity.

2.3.2: Concordance with DepMap. Since there are core fitness genes shared by all cells, we expect gene scores should roughly correlate with average DepMap dependency scores.

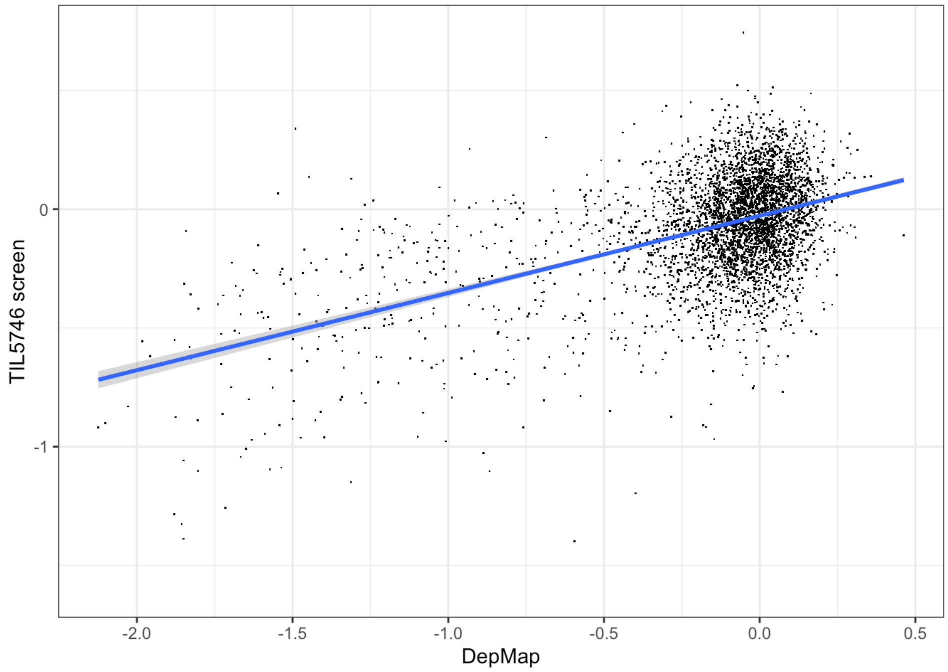


The gene scores for this screen recapitulate the pattern of core essential genes from DepMap.

**Supplemental Data 2:** OT1 / B16-Ova screen QC

Background: This screen investigated the effect of CRISPR knockout on OT1 T cells in vivo. Two sub-genome libraries, lib30 and lib31 were used.

3. Results: The screen analysis was performed on zorya. Both libraries (lib30 and lib31) were merged together to perform the screen analysis, with normalized clone counts and a merged sgRNA library file. A QC report was generated with clean_count output saved.

   1.1: Description of samples included in analysis:

| Library | Day of Analysis | Number of Samples |
|---|---|---|
| Lib30 | Day 14 | 7 |
| Lib30 | Day 21 | 7 |
| Lib31 | Day 14 | 7 |
| Lib31 | Day 21 | 7 |

4. Library/Screen QC

2.1: Screen sequencing purity: What fraction of sequenced reads match to the expected library design?

Recovered sgRNAs: 97%. Missing: 3%.

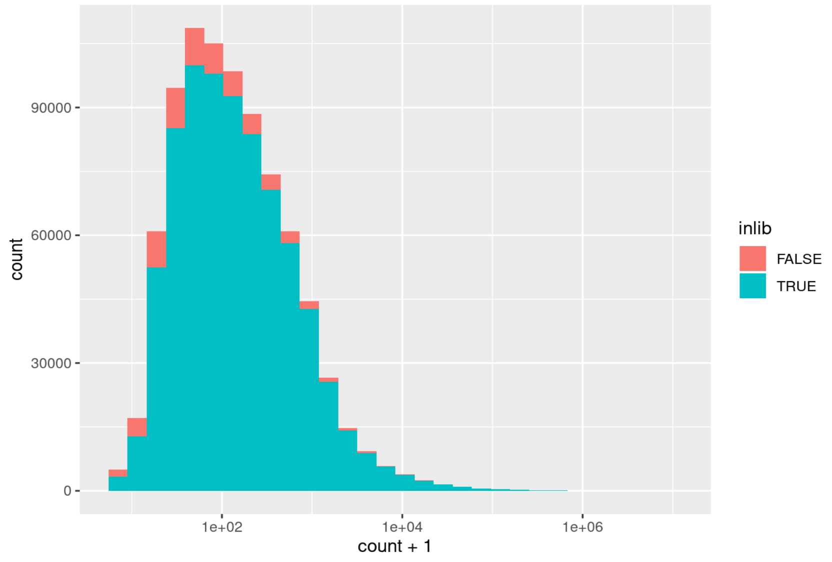


The majority of reads match to the library design, and the sequences matching the library design have much higher counts than sequences that are not part of the library design.

2.2: Screen Coverage

We expect the majority of the sgRNA library to be recovered.

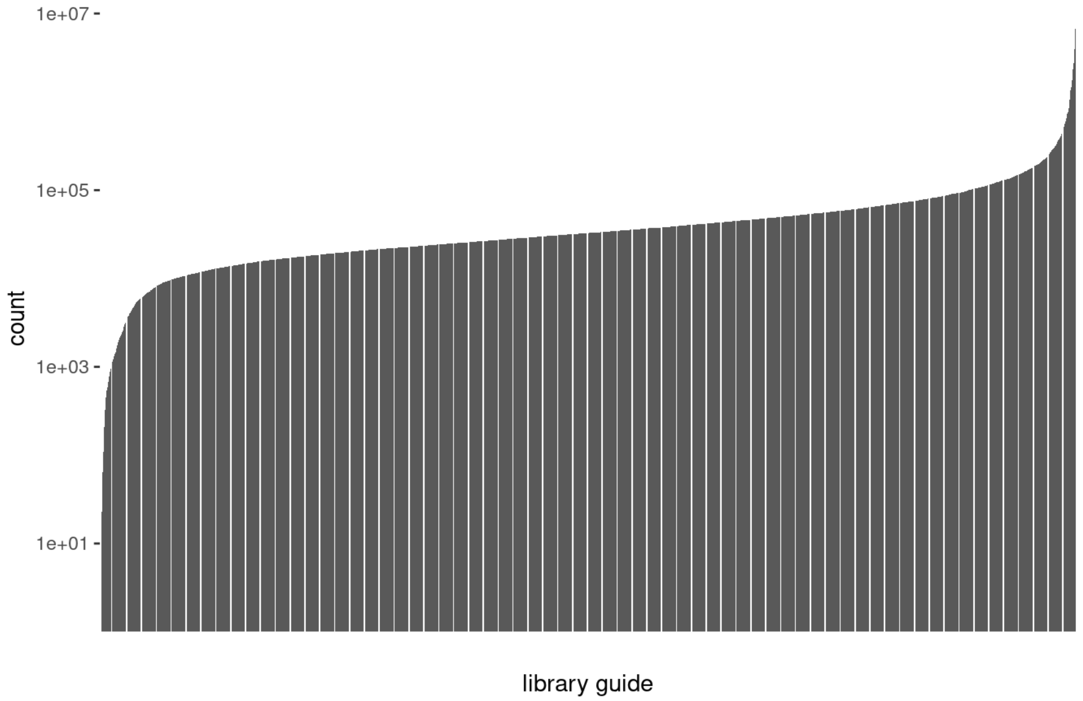


Majority of guides have uniform counts.

2.3: Gene Level QC

2.3.1: Negative controls should drop out

lib30 and lib31 included controls targeting multiple positions across the genome (type="control_multicutter"). We expect these guides to drop out compared to the average guide in the library.

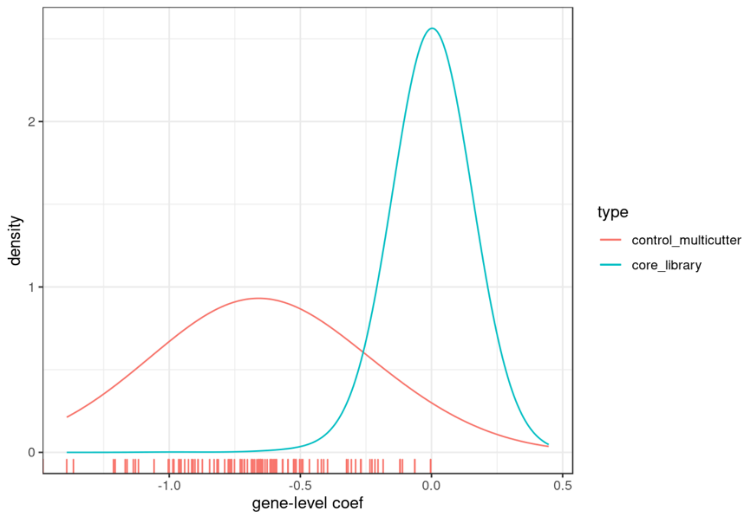


Control guides targeting high-copy regions of the genome drop out, confirming Cas9 activity.

2.3.2: Neutral controls should not enrich/deplete. Two other types of control guides were included in the library:

- **control_ORuniquecutter** are guides cutting unique locations within Olfactory receptor genes, which are not expressed in T cells.
- **control_randomnoncutting** guides don't target any sequence in the mouse reference genome and are not expected to cut. Non-cutting guides are expected to have a slight growth advantage compared to guides that create a double stranded break.

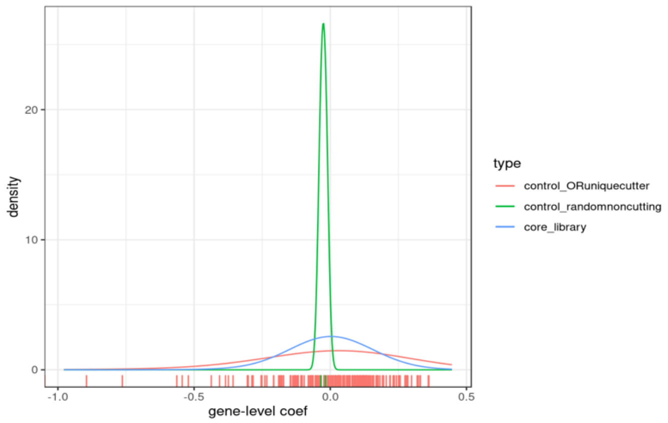


Both Olfr cutters and noncutting guides distributed nicely with coefs centered at 0, with no growth advantage observed for random-noncutting guides.

**Supplemental Data 3:** PMEL / MC38-gp100 screen QC

Background: This screen investigated the effect of CRISPR knockout on PMEL T cells in vivo. Two sub-genome libraries, lib30 and lib31 were used.

5. Results: The screen analysis was performed on zorya. Both libraries (lib30 and lib31) were merged together to perform the screen analysis, with normalized clone counts and a merged sgRNA library file. A QC report was generated with clean_count output saved.

   1.1: Description of samples included in analysis:

| **Library** | **Day of Analysis** | **Number of Samples** |
|---|---|---|
| Lib30 | Day 14 | 7 |
| Lib31 | Day 14 | 7 |

6. Library/Screen QC

2.1: Screen sequencing purity: What fraction of sequenced reads match to the expected library design?

Recovered sgRNAs: 82%. Missing: 18%.

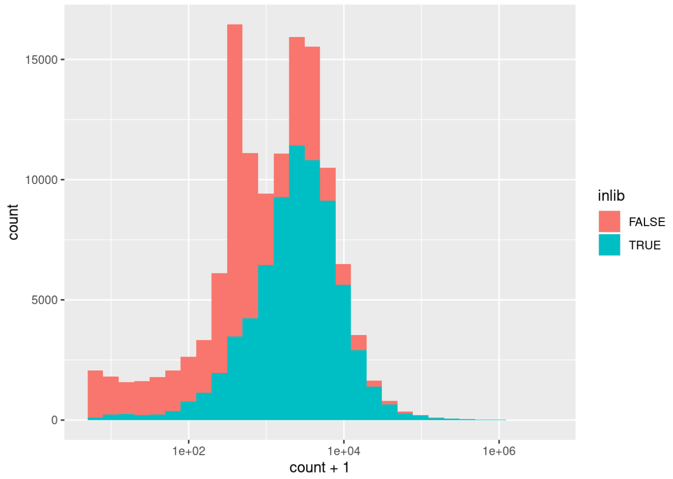


The majority of reads with high counts match to the library design. For guide sequences with counts below 1,000, majority of them are noise sequences that doesn't belong to the library sgRNA.

2.2: Screen Coverage

We expect the majority of the sgRNA library to be recovered.

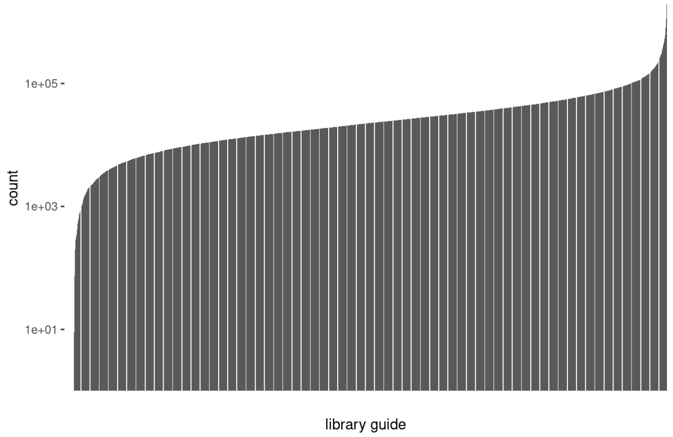


Majority of guides have uniform counts.

2.3: Gene Level QC

2.3.1: Negative controls should drop out

lib30 and lib31 included controls targeting multiple positions across the genome (type="control_multicutter"). We expect these guides to drop out compared to the average guide in the library.

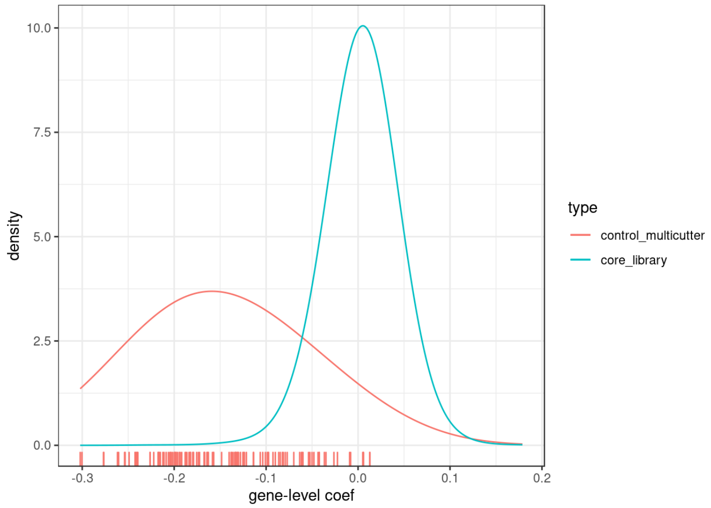


Control guides targeting high-copy regions of the genome drop out, confirming Cas9 activity.

2.3.2: Neutral controls should not enrich/deplete. Two other types of control guides were included in the library:

- **control_ORuniquecutter** are guides cutting unique locations within Olfactory receptor genes, which are not expressed in T cells.
- **control_randomnoncutting** guides won't target a specific genomic location. They should have growth advantage.

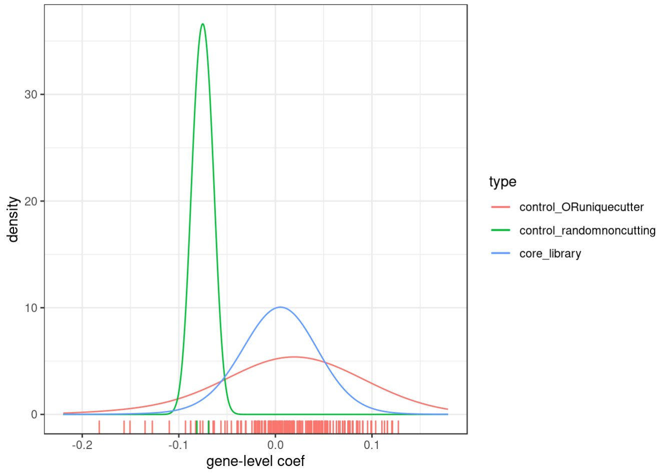


Olfr cutters distributed as expected, with coefs centered at 0. Non-cutting guides are limited in numbers; Using Olfr cutters as reference, both non-cutting guides stay within the dynamic range of neutral guides. No growth advantage observed for random-noncutting guides.

**Supplemental Figures**

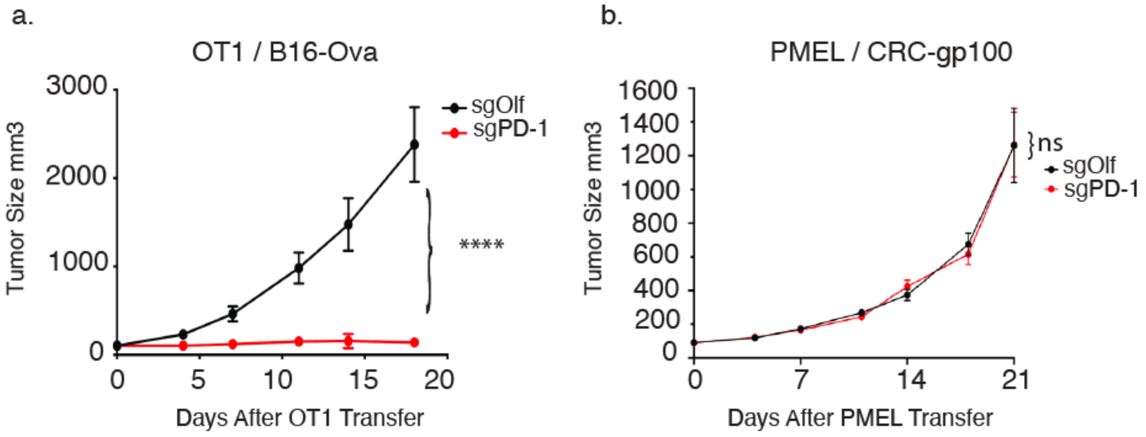


**Figure S1: Development of syngeneic tumor models sensitive and insensitive to inactivation of PD-1 in adoptively transferred TCR-Tg CD8 T cells**

C57BL/6 mice bearing the indicated tumor at a median size of $100mm^3$ on the flank were treated with $3x10^6$ OT1 CD8 T cells or $7x10^6$ PMEL CD8 T cells inactivated for either PD-1 (sgPD-1) or OLF1 (sgOlf) as indicated. **(a)** B16-Ova growth curves over time following transfer demonstrates that sgPD-1 OT1 T cells can control growth **(b)** MC38-gp100 growth curves over time following transfer demonstrates that sgPD-1 PMEL CD8 T cells are unable to control tumor growth. Data expressed as mean +/- SEM, with ns = no significance and **** = p value < 0.0001 by 2-way ANOVA between the indicated comparator groups.

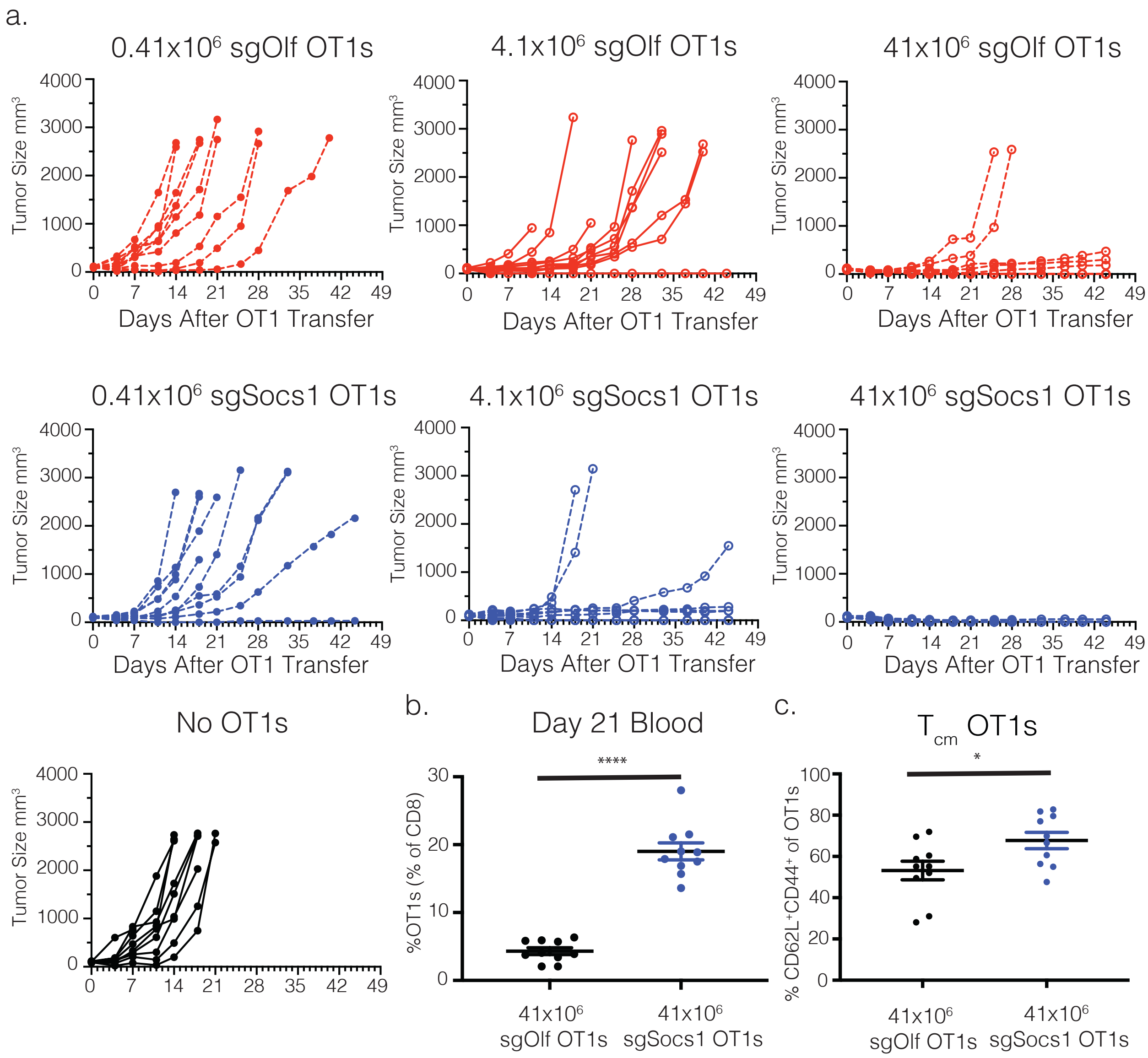


**Figure S2: Inactivation of SOCS1 enhances the in vivo anti-tumor potency of CD8 T cells and drives the accumulation of $CD44^+CD62L^+$ $T_{cm}$ memory cells in blood**

C57BL/6 mice bearing B16-Ova tumors cells at a median size of 100mm$^3$ on the flank were treated with SOCS1 (sgSocs1) or OLF1 (sgOlf) OT1 CD8 T cells as indicated. Editing efficiencies for target genes were 91% for sgSocs1 and 71% for sgOlf. **(a)** Tumor growth curves over time are depicted. **(b)** The frequency of $Va2^+Vb5.1^+$ OT1s present in the peripheral blood of mice receiving a dose of 41x10$^6$ OT1s is depicted 21 days following transfer. **(c)** the frequency of $CD44^+CD62L^+$ $T_{cm}$ and $CD44^+CD62L^-$ $T_{em}$ OT1s were quantified in mice receiving 41x10$^6$ OT1s 21 days following transfer. Each dot represents an individual mouse, with **** = p value < 0.0001; and * = p value < 0.05 by Student's t test between the indicated comparator groups.

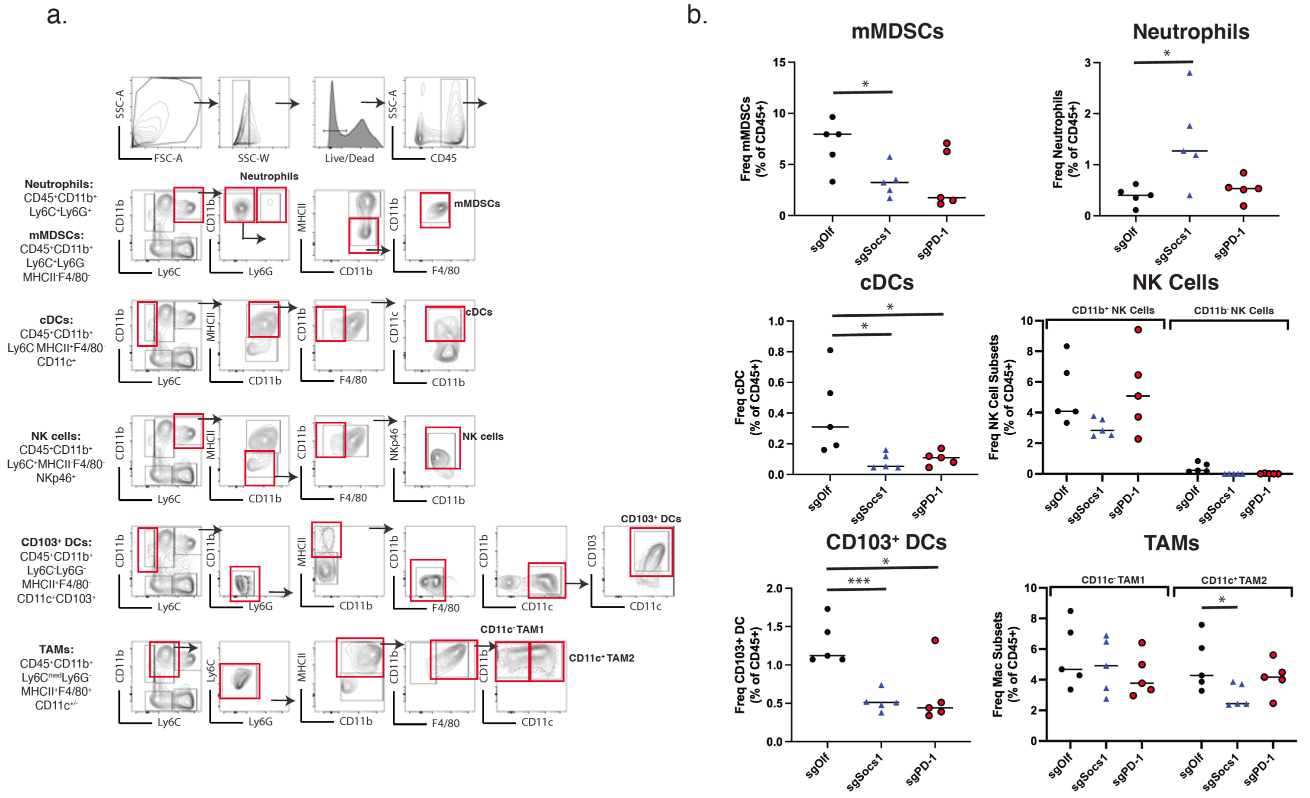


**Figure S3: Inactivation of SOCS1 in transferred CD8 T cells drives increased neutrophils and reduced mMDSCs, CD103⁺ DCs and TAM2s in the TME**

C57BL/6 mice bearing B16-Ova tumors cells at a median size of 100mm$^3$ on the flank were treated with SOCS1 (sgSocs1) or OLF1 (sgOlf) OT1 CD8 T cells as indicated. Tumors were harvested at Day 7, and FACS analyses performed to quantify innate cell populations. **(a)** FACS gating scheme is depicted to identify neutrophils, monocytic myeloid-derived suppressor cells (mMDCs), classical dendritic cells (cDCs), natural killer cells (NK cells), CD103+ dendritic cells (CD103$^+$ DCs), and type 1 and 2 tumor-associated macrophages (CD11c$^-$ TAM1, CD11c$^+$ TAM2). . **(b)** The frequency of each innate cell population is depicted by treatment group, with each symbol representing an individual mouse, with **** = p value < 0.0001; and * = p value < 0.05 by Student's t test between the indicated comparator groups.

a.

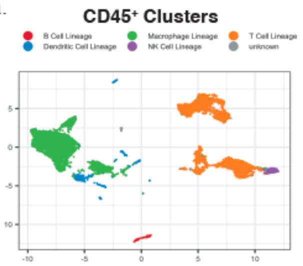


b.

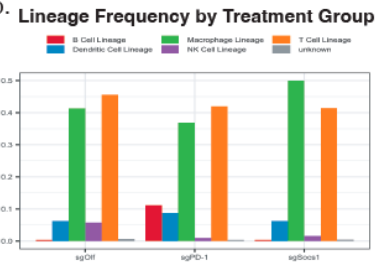


c.

T cell Cluster Total Counts

| | C0 | C1 | C2 | C3 | C4 | C5 | C6 | C7 | C8 | C9 | C10 | C11 | Total |
|---|---|---|---|---|---|---|---|---|---|---|---|---|---|
| cell counts | 2000 | 1155 | 1084 | 812 | 713 | 639 | 358 | 356 | 348 | 315 | 227 | 116 | 8123 |
| % sgSocs1 | 627 | 500 | 360 | 407 | 260 | 433 | 14 | 138 | 90 | 59 | 51 | 44 | |
| % sgPD-1 | 211 | 139 | 255 | 192 | 138 | 109 | 311 | 49 | 127 | 63 | 49 | 46 | |
| %sgOlf | 1162 | 516 | 469 | 213 | 315 | 97 | 33 | 169 | 131 | 315 | 127 | 26 | |
| Totals | 2000 | 1155 | 1084 | 812 | 713 | 639 | 358 | 356 | 348 | 315 | 227 | 116 | |

d.

T cell Cluster Marker Genes

| | | | | | | | | | | |
|---|---|---|---|---|---|---|---|---|---|---|
| C0 | Tyrobp | Fcer1g | Klra7 | Cd7 | Pmepa1 | Emb | Pde2a | Klra1 | Trbv12-2 | 1700025G04Rik |
| C1 | Ccl5 | Gzmk | Lgals3 | Trbv31 | Trbv14 | S100a4 | S100a6 | Crgln1 | Rgs16 | Nkg7 |
| C2 | Trbv12-1 | Ccr7 | 2310001H17Rik | Trbc2 | Hspb1 | Dapl1 | Fam162a | Slamf6 | Dusp4 | Gdl2 |
| C3 | Hist1h2ae | Hist1h1b | Ube2c | Pclaf | Top2a | Mki67 | Tuba1b | Hist1h2ap | Tubb5 | Stmn1 |
| C4 | Ramp3 | Areg | Tnfsf11 | Rora | Cd4 | Cd40lg | Odc1 | Gatl | Icos | Tmem64 |
| C5 | Gm156 | Gzmb | Ccl3 | Ccl4 | Mt1 | Plac8 | Spp1 | Lgals3 | Serpinb1a | S100a4 |
| C6 | Hspa1b | Hspa1a | Igfbp4 | Fos | Klf2 | Jun | Dapl1 | Gm20400 | Dnajb1 | Lef1 |
| C7 | Xcl1 | Ccl1 | Pclaf | Top2a | Tuba1b | Hmgb2 | Stmn1 | Ube2c | Mki67 | Ran |
| C8 | Ccr7 | Fam241a | Tubb2a | Rflnb | Lef1 | Satb1 | Gbp2 | Igfbp4 | Dusp10 | Igtp |
| C9 | Hspa1b | Hspa1a | Jun | Fos | Klf2 | Dnajb1 | Id3 | Dusp1 | Hspb1 | Cd69 |
| C10 | Foxp3 | Tnfrsf4 | Ctla4 | Areg | Maf | Izumo1r | Capg | Lrrc32 | Il2ra | Sdf4 |
| C11 | Gm26917 | Lars2 | Gm42418 | Ccnb1ip1 | AY036118 | Malat1 | Cacnb3 | Gm10800 | Mreg | Gm49774 |

e.

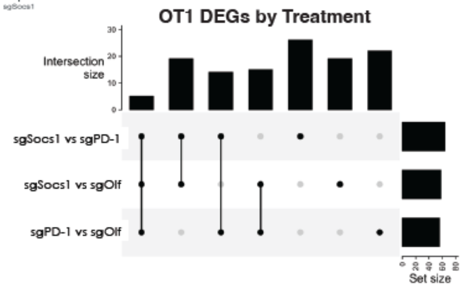


f.

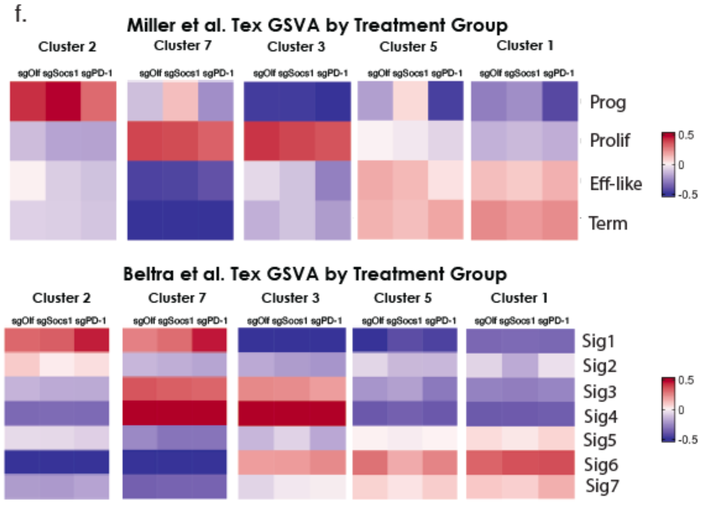


g.

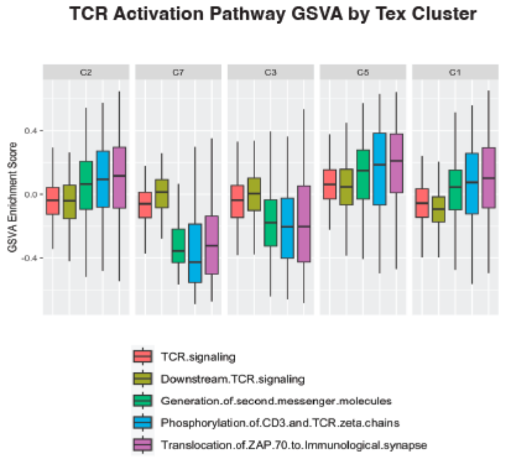


h.

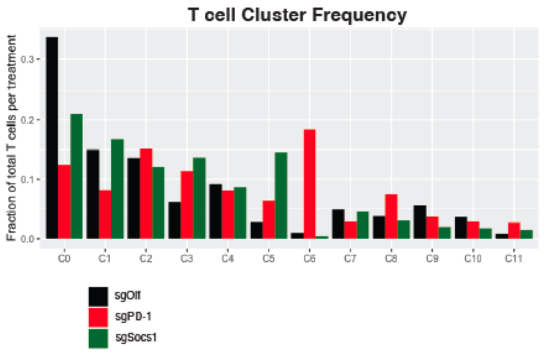


i.

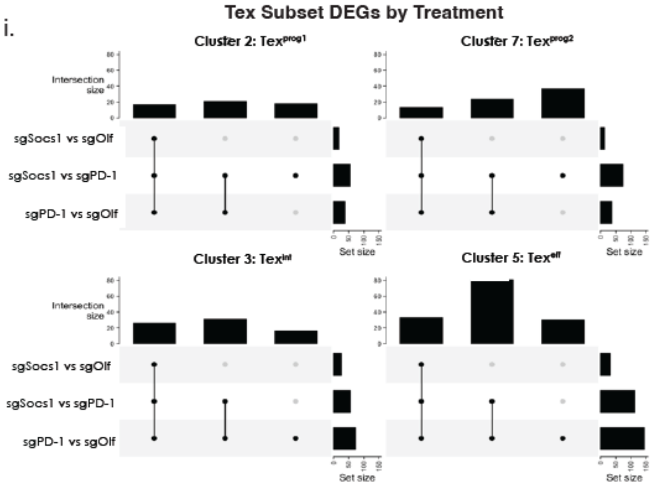

Tex Subset DEGs by Treatment

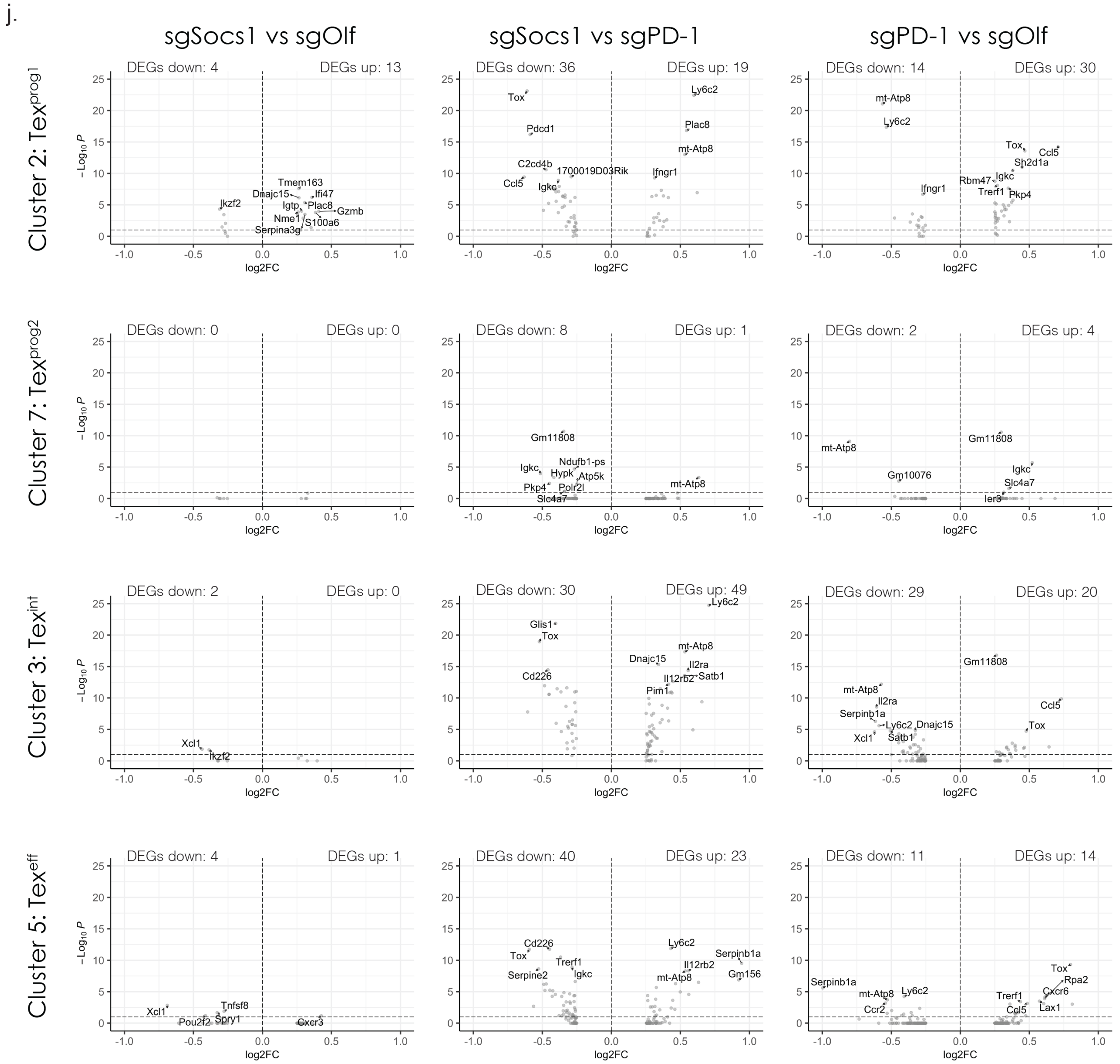


**Figure S4:**

**scRNA-Seq of sgOlf, sgSocs1 and sgPD-1 OT1s from the TME 7 days following transfer. (a)** Distribution of general lymphocytes populations from CD45⁺ cells from the TME. Cells were annotated computationally, with the Haemosphere mouse RNA-Seq database(3) used to verify the predominate lymphocyte clusters. **(b)** General lymphocyte population frequency per treatment group. **(c)** Cluster-wise cell counts per treatment group. **(d)** Top 10 DEGs (by avgLog2FC) per cluster. Significance was determined using the Wilcoxon Rank Sum test and genes with an adjusted p value < 0.1 were considered significant. **(e)** UpSet plot of DEG counts between treatment groups within the OT1 population. **(f)** Heatmap of median GSVA scores of

Miller et al.(3) Tex gene signatures and Beltra et al.(4) Tex gene signatures per Tex cluster by treatment group. **(g)** Boxplot of GSVA scores of five TCR activation-related gene signatures from Reactome by Tex cluster. **(h)** Barplot of T cell cluster frequency by treatment group. **(i)** UpSet plots of DEG counts between treatment groups per Tex cluster. **(j)** Volcano plots depicting DEGs between treatment groups per Tex cluster within the OT1 population. Names of the top 10 genes (by adjusted p value) are highlighted for each comparison. If fewer than 10 genes have an adjusted p value $< 0.1$ for a comparison, only the names of genes with an adjusted p value $< 0.1$ are highlighted.

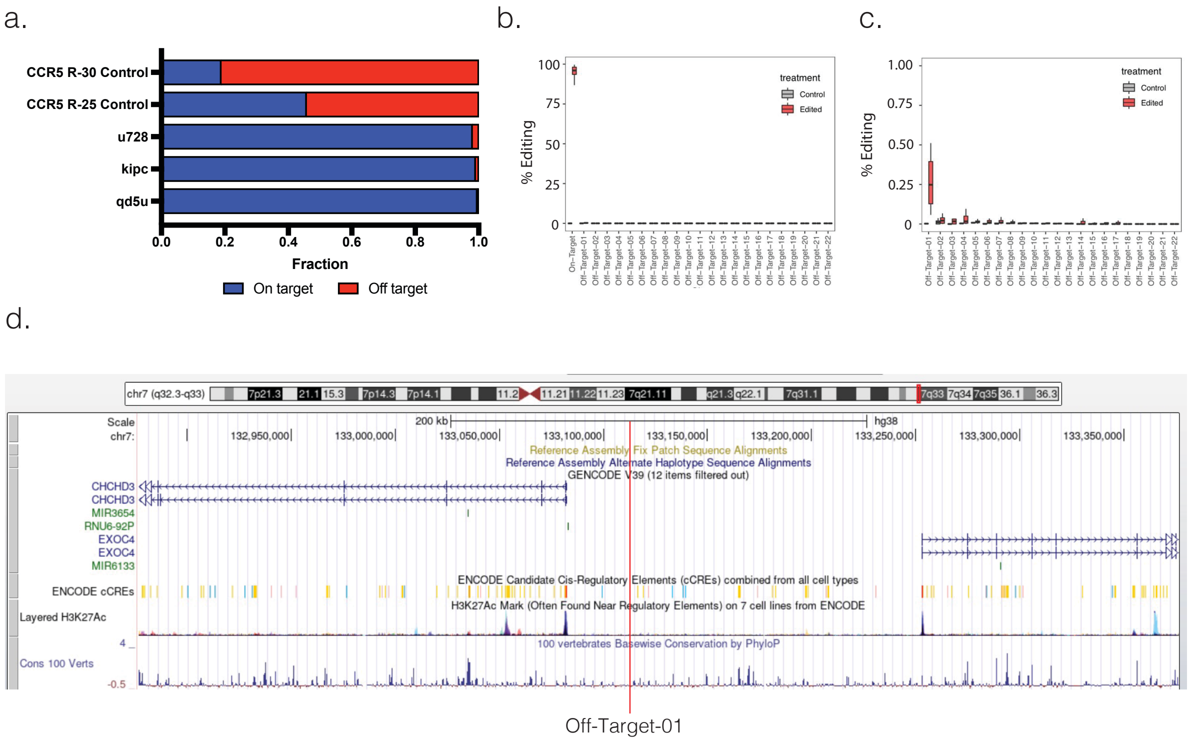


**Figure S5: Identification of the genomic locations of u728 sgRNA off-target cut sites**
**(a)** GUIDE-Seq was used to quantify and characterize the frequency of on- and off-target cut sites in primary human T cells. Two sgRNAs targeting CCR5 (R-30 and R-25) with high off-target editing frequencies were included as benchmarks, with the SOCS1 on-target editing efficiency depicted in blue, and the cumulative editing efficiency of off-target loci depicted in red for the u728, kipc and qd5u sgRNAs. **(b)** Amp-Seq confirmation of on- and off-target editing for the u728 sgRNA in human TIL including 22 potential off-target sites identified in (a). The u728 sgRNA achieved a median of 96% editing of the SOCS1 gene. Off-Target-01 confirmed as reaching statistically significant levels of 0.3%. **(d)** The genomic location of the Off-Target-01 site is in an intragenic region on Chromosome 7 with no known regulatory regions.

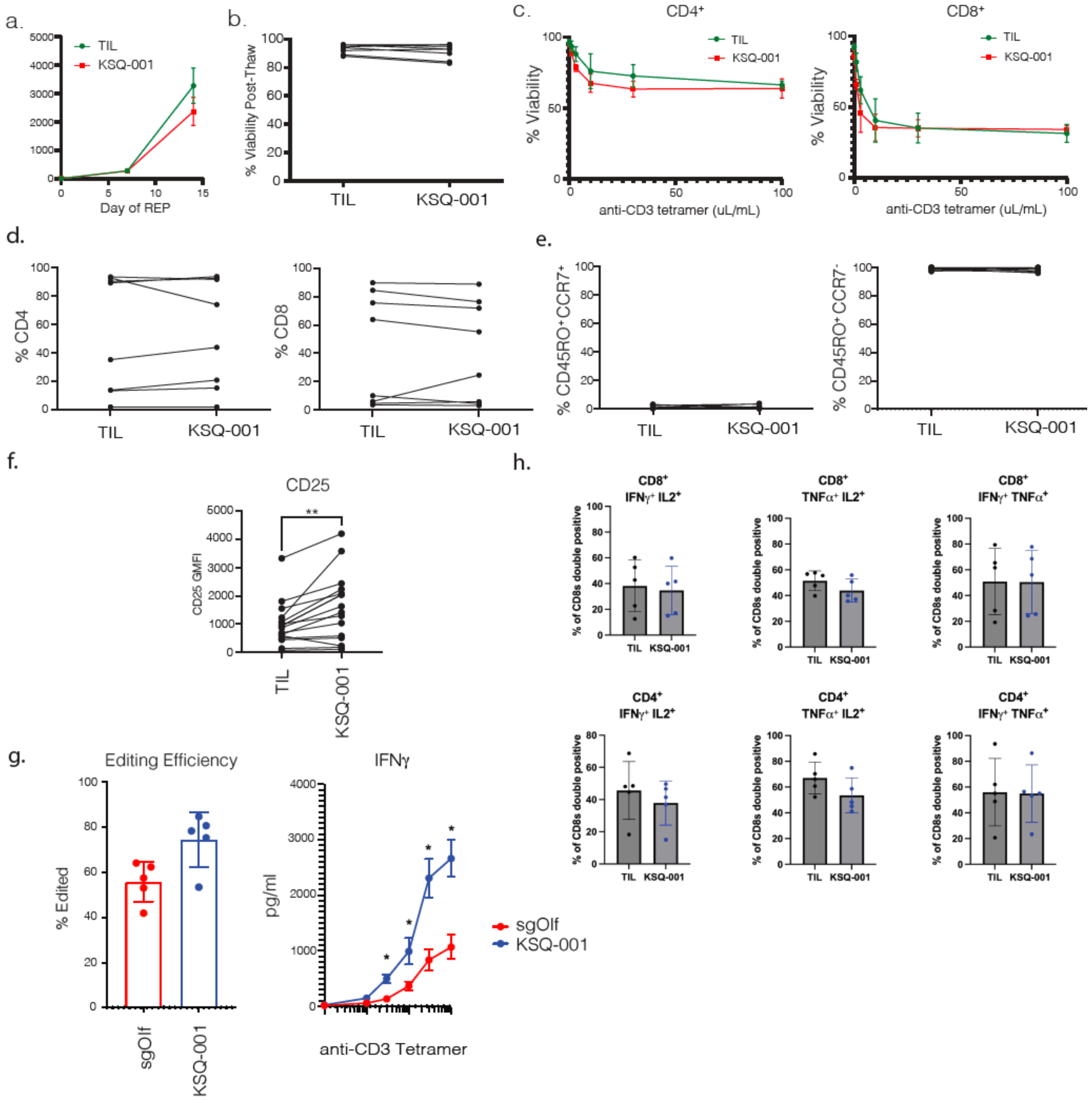


**Figure S6: Characterization of KSQ-001**

TIL and KSQ-001 were manufactured. (**a**) Fold expansion of TIL and KSQ-001 cells over 14 days of REP. (**b**) Following manufacture and cryopreservation, TIL and KSQ-001 were thawed and viability assessed by AOPI staining, with each dot reflecting an independent paired donor (**c**) To evaluate the ability of KSQ-001 to undergo activation-induced cell death (AICD) following activation, cryopreserved TIL and KSQ-001 cells were thawed, rested overnight, and activated with the indicated concentration of CD3 tetramer for 18 hours. The AICD profile of CD4 and CD8 within TIL and KSQ-001 was measured by FACS analysis of Caspase-3 staining. (**d**) The frequency of donor-paired CD4 and CD8 cells within KSQ-001 and TIL is depicted. Each dot reflects an independent donor. (**e**) The frequency of central memory cells (CCR7$^+$, CD45RO$^+$) and effector memory cells ($CCR7^-$, $CD45RO^+$) in TIL and KSQ-001 following manufacture are displayed. **(f)** CD25 expression of TIL and KSQ-001 across independent donors. **(g)** TIL and KSQ-001 were manufactured with the same donor, with TIL undergoing electroporation of sgRNA/Cas9 RNPs targeting an Olfactory gene (sgOlf). Editing efficiency of the Olfactory and SOCS1 genes are depicted in the graph on left, and IFNg production following activation with anti-CD3 tetramers at the indicated concentration is shown on the right. **(g)** Polyfunctionality of TIL and KSQ-001 from five independent donors was assessed, with frequency of CD8 or CD4 T cells producing multiple cytokines by intracellular cytokine stain (ICS) assessed as indicated. Each dot reflects an independent donor, with ** = p value <0.01 and * = < 0.05 by a Student's *t* test between the indicated comparator groups.

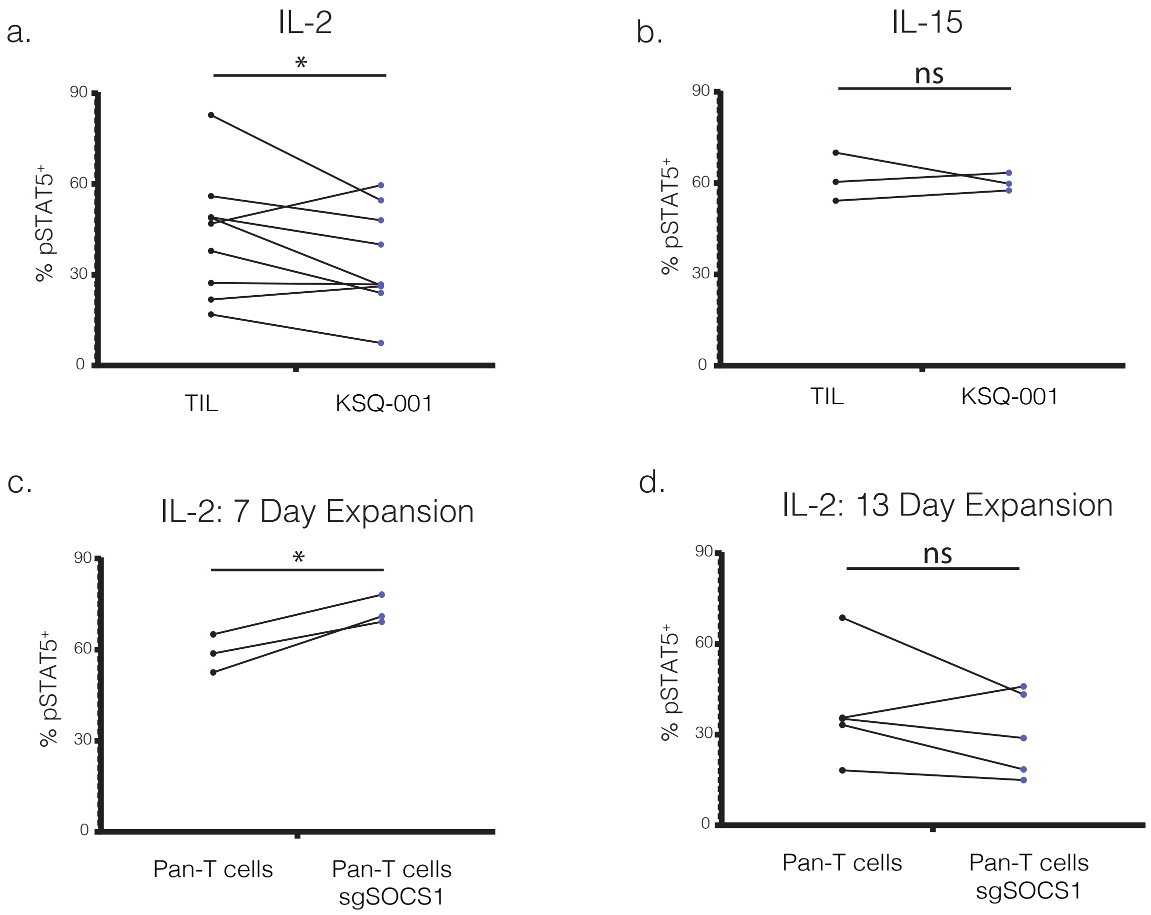


**Figure S7: Sensitivity of KSQ-001, TIL, and sgSOCS1-edited CD3$^{+}$ T cells to IL-2 and IL-15**
The sensitivity of KSQ-001 and TIL to IL-2 and IL-15 through STAT5 phosphorylation (pSTAT5) was evaluated. Following REP and cryopreservation, donor-paired TIL and KSQ-001 were thawed, rested, and activated with the indicated cytokine. **(a)** pSTAT5 signals between TIL and KSQ-001 following IL-2 activation. **(b)** pSTAT5 signals between TIL and KSQ-001 following IL-15 activation. **(c)** CD3$^{+}$ Pan-T cells were activated and edited sgSOCS1, with sensitivity to IL-2 through pSTAT signals evaluated at 7 days following activation in comparison to unengineered cells. **(d)** Same experiment as in (c), but with pSTAT5 sensitivity to IL-2 evaluated at 13 days following initial activation. Statistical significance: * = $p < 0.05$ using a Student's *t* test between the indicated groups. ns = not significant
